# Intertwined signatures of desiccation and drought tolerance in grasses

**DOI:** 10.1101/662379

**Authors:** Jeremy Pardo, Ching Man Wai, Hannah Chay, Christine F. Madden, Henk W.M. Hilhorst, Jill M. Farrant, Robert VanBuren

## Abstract

Grasses are among the most resilient plants and some can survive prolonged desiccation in semi-arid regions with seasonal rainfall. This vegetative desiccation tolerance has arisen independently multiple times within the grass family, but the genetic elements that differentiate desiccation tolerant and sensitive grasses are largely unknown. Here we leveraged comparative genomic approaches with the resurrection grass *Eragrostis nindensis* and the closely related desiccation sensitive cereal *Eragrostis tef* to identify changes underlying desiccation tolerance. We extended the analyses to include the grasses maize, sorghum, rice, and the model desiccation tolerant grass *Oropetium thomaeum* to identify broader evolutionary conservation and divergence. We identified changes in chromatin architecture and expression dynamics related to desiccation in *E. nindensis*. It was previously hypothesized that transcriptional re-wiring of seed desiccation pathways confers vegetative desiccation tolerance. We demonstrate that the majority of seed dehydration related genes show similar expression patterns in leaves of desiccation tolerant and sensitive species during dehydration. However, we discovered a small set of orthologs with expression specific to leaves of desiccation tolerant species, and seeds of sensitive species. This supports a nuanced role of seed-related genes where many overlap with typical drought responses but some crucial genes are desiccation specific in resurrection plants.

## Introduction

Approximately 470 million years ago charophyte green algae emerged from their watery habitat to colonize land (Delwiche and Cooper, 2015). Exposure to a harsh dry atmosphere was the main biophysical constraint facing early land plants, resulting in strong selective pressure favoring adaptive mechanisms to prevent dehydration (Bateman et al., 1998). The evolution of desiccation tolerant seeds and pollen was critical to the success of seed plants and the majority of extant lineages retain both abilities (Franchi et al., 2011). In contrast, vegetative desiccation tolerance, while widespread among plant lineages, is a relatively uncommon trait (Oliver et al., 2000). The appearance of vegetative desiccation tolerance in phylogenetically distant lineages suggests multiple independent evolutionary origins. In the ecologically and economically important plant family Poaceae, vegetative desiccation tolerance likely evolved independently a minimum of nine times (Gaff and Oliver, 2013). The current consensus hypothesis is that vegetative desiccation tolerance in angiosperms arose convergently through re-wiring of common seed desiccation pathways (Costa et al., 2017; VanBuren et al., 2017).

Transcriptomic studies with desiccation tolerant angiosperms consistently show activation of normally seed specific genes under drought conditions (Costa et al., 2017; Gechev et al., 2013; Rodriguez et al., 2010; VanBuren et al., 2018a, 2017; Yobi et al., 2017; Zhu et al., 2015). However, many of these genes are also highly expressed during drought response in desiccation sensitive species. The phytohormone abscisic acid (ABA) is critical for seed maturation and drought responses, and is hypothesized to play a major role in regulating desiccation tolerance (Gaff and Oliver, 2013; Manfre et al., 2009; Shinozaki and Yamaguchi-Shinozaki, 2007). Thus, many of the downstream genes that are activated via ABA dependent mechanisms are expressed broadly during seed development and drought as well as under desiccated conditions in resurrection plants. Accumulation of osmoprotectants and activation of reactive oxygen species quenching mechanisms are also shared responses between these three conditions. Thus, it is important to distinguish between desiccation tolerance specific responses from general drought responses in order to uncover the genetic architecture of desiccation tolerance.

Identification of genes that are expressed in leaves of desiccation tolerant, but not desiccation sensitive plants during drought could provide targets to improve drought tolerance in crops. While numerous transcriptomic studies of desiccation tolerant plants have been published, few previous studies have compared the responses of desiccation sensitive and desiccation tolerant plants with a close phylogenetic relationship. Previous work comparing the eudicot species *Lindernia brevidens* (desiccation tolerant) and *Lindernia subracemosa* (desiccation sensitive) provided some insight into genes that are involved in desiccation tolerance and not drought (VanBuren et al., 2018a). However, the Linderniaceae family is of little economic importance and is only distantly related to any crop plants making it difficult to translate these discoveries. Cereals from the grass family (Poaceae), are the most important crops for global food security (Food and Agriculture Organization of the United Nations., 2012), and work with desiccation tolerant Poaceae species is likely more readily translatable. The Chloridoideae subfamily of grasses contains the majority of desiccation tolerant species with at least 6 independent phylogenetic origins. Chloridoideae also contains the cereals finger millet (*Eleusine coracana*) and tef (*Eragrostis tef*), which are widely consumed in semi-arid regions of Eastern Africa and Asia. To our knowledge, *Eragrostis* is the only genus with both desiccation tolerant, and crop plant species. Thus, *Eragrostis* and the Chloridoideae subfamily more broadly, is an ideal system to identify genes involved in desiccation tolerance that are potential targets for improving drought resilience in crops.

Chromosome-scale genome assemblies of the Chloridoideae grasses *Eragrostis tef* (desiccation sensitive) and *Oropetium thomaeum* (desiccation tolerant) were recently completed (VanBuren et al., 2019b, 2018b). Here, we sequenced the desiccation tolerant grass *Eragrostis nindensis* and performed detailed comparative genomics within Chloridoideae and across the grasses to search for patterns of convergence in the evolution of desiccation tolerance. We conducted parallel dehydration experiments with *E. nindensis* and *E. tef* using matched physiological sampling points to identify signatures that distinguish drought and desiccation responses. We also leveraged seed expression data of *E. tef* and other grass species to test if desiccation tolerance in grasses arose through seed pathway re-wiring. Together, our results identified a series of genomic features and expression changes unique to desiccation tolerant grasses. We also propose a new model that re-wiring of seed pathways is a common feature of desiccation tolerance and more general water stress responses, with only a few pathways uniquely expressed in seeds, and leaves of desiccation tolerant plants.

## Results

### Assembly and annotation of the E. nindensis genome

Comparative systems with phylogenetically similar desiccation sensitive and tolerant species are a powerful tool to elucidate the genetic basis of desiccation tolerance. Only one previous study has conducted genome-wide comparisons between a desiccation sensitive and tolerant angiosperm (VanBuren et al., 2018a), and no such systems are currently available for the grasses. We assembled a draft genome of the desiccation tolerant grass *E. nindensis* and compared it to the recently sequenced *E. tef* genome to identify genetic elements associated with desiccation tolerance. We utilized a single molecule real-time sequencing approach to overcome assembly issues related to tetraploidy and heterozygosity in *E. nindensis*. In total, we generated 64 Gb of PacBio data representing 63x coverage of the 1.0 Gb *E. nindensis* genome. PacBio reads were error corrected and assembled using Canu, followed by polishing with Pilon using high-coverage Illumina data. Canu parameters were optimized to accurately assembly all haplotypes, yielding an initial *E. nindensis* genome assembly with 16,706 contigs spanning 1.96 Gb, or roughly twice the haploid genome size and a contig N50 of 220 kb (Supplemental Table 1). We utilized the Pseudohaploid algorithm to filter out redundant haplotypes from the assembly (see methods for details). This filtering approach yielded a total haploid assembly of 986 Mb across 4,368 contigs with an N50 of 520kb. This assembly is referred to as *E. nindensis* V2.1.

We annotated 116,452 genes in the *E. nindensis* genome using the MAKER-P pipeline (Campbell et al., 2014). Of these, 79,755 were syntenic with *E. tef*, and 80,997 had at least one ortholog, syntenic or otherwise in *E. tef*. 84,603 genes have at least one pfam domain. Overall, 98,294 genes (84.4%) were orthologous to *E. tef* or contained pfam domains. We combined this set of genes with the 58,602 genes with detectable expression (defined as ∑*TPM* > 1 across all conditions sampled) to create a set of 107,683 “high confidence” gene models. Of these high confidence genes, 74.1% were syntenic with *E. tef*, and 80.0% of genes with detectable expression were syntenic, suggesting sufficient collinearity for genome wide comparisons. We used the Embryophyta Benchmarking Universal Single-Copy Orthologs (BUSCO) to evaluate the completeness of our annotation. We found copies of most of the 1440 Embryophyta BUSCOs (92.1% complete, 95.6% complete or fragmented), and the majority were duplicated (65.6%; 946), which is consistent with the polyploid nature of *E. nindensis*.

Similar to ~90% of Chloridoid grasses, *E. nindensis* is a complex polyploid. Using DAPI stained root tips, we identified 40 chromosomes, which matches the karyotype of allotetraploid *E. tef* (2n=4x=40) and prior karyotyping of *E. nindensis* (Supplementary Figure 1). The *E. nindensis*, *E. tef*, and *O. thomaeum* genomes are highly syntenic, with conserved gene content and order (Figure 1). A high proportion of the *E. tef* and *E. nindensis* genomes matched the expected 2:2 ratio of syntenic gene blocks given their shared polyploidy, but a significant portion of the *E. nindensis* genome have 3 or 4 blocks for each homeologous region of *E. tef* (Supplemental Figure 2b). *E. nindensi*s and the diploid *O. thomeaum* have 2:1 synteny with similarly duplicated blocks of 3 or more in some regions (Supplemental Figure 2a). This syntenic pattern is likely a result of assembling multiple haplotypes for each homeologous region in *E. nindensis* and a single haplotype for *E. tef* and *O. thomaeum*. This is supported by the distribution of synonymous substitutions (Ks) between orthologous genes in *E. nindensis* (Supplemental Figure 3). We observed a strong peak of Ks corresponds to haplotype sequences and homeologous gene pairs. The high degree of collinearity and shared gene content between the three sequenced Chloridoideae grasses allowed us to identify shared and unique genomic signatures of desiccation tolerance.

**Figure 1.**
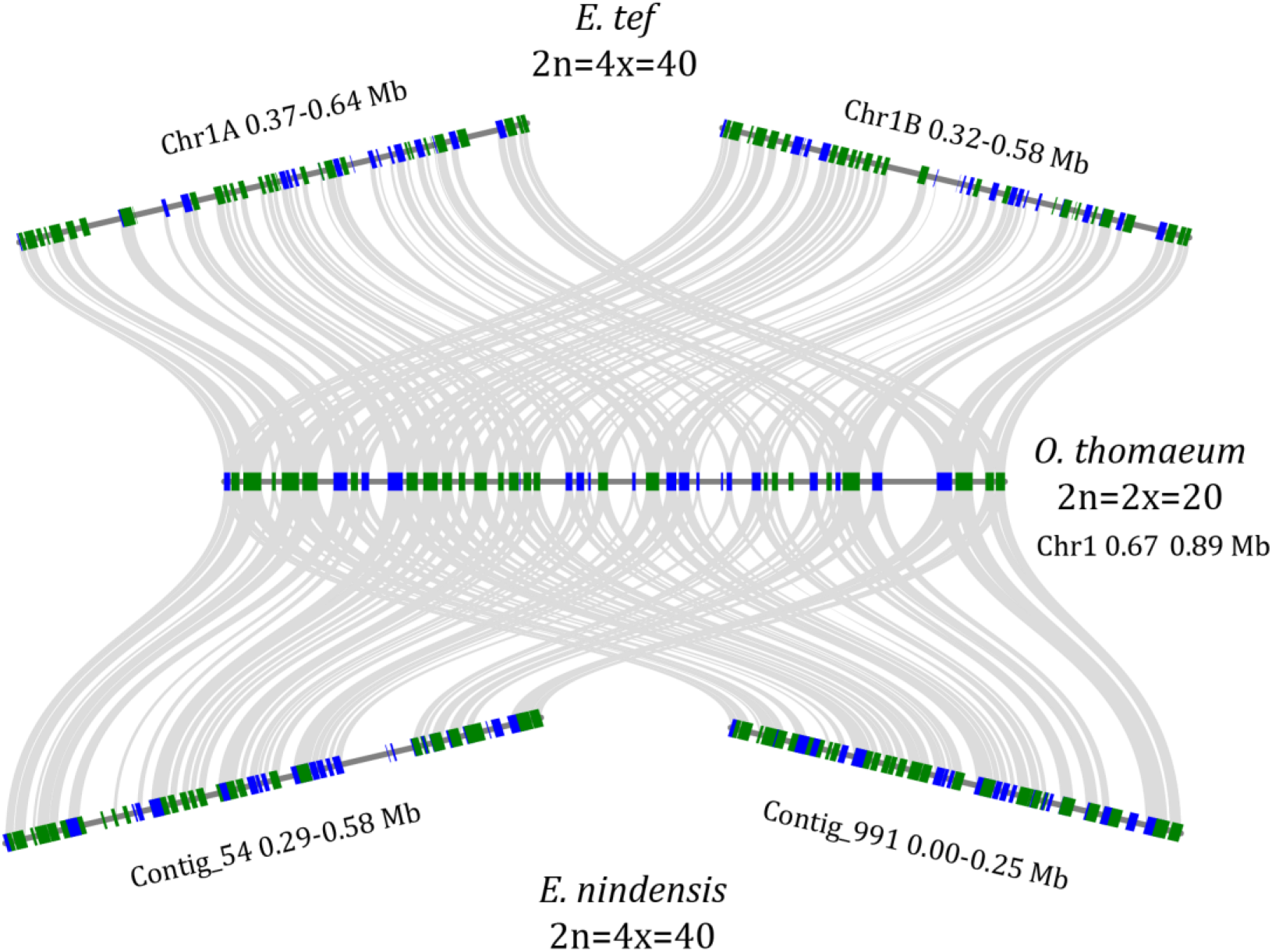
Collinearity of Chloridoideae grasses. Microsynteny of collinear regions between allotetraploid *E. nindensis* and *E. tef* and diploid *O. thomaeum* is shown. Genes in the forward and reverse orientation are shown in green and blue respectively, and syntenic gene pairs are connected by grey lines.

### Comparative water deficit responses between E. nindensis and E. tef

We sampled parallel timepoints across dehydration timecourses in *E. tef* and *E. nindensis* to distinguishing drought and desiccation associated expression patterns (Figure 2a). *E. tef* and *E. nindensis* utilize water at different rates, so we used relative water content (RWC) as a proxy of drought stress severity for sample comparison. At the start of the drought experiment (0 Zeitgeber time, ZT day 1) the mean RWC of the *E. tef* samples was 75.8% while the RWC of *E. nindensis* samples was 60.4% (Figure 1b). By 8 ZT on day 1 (D1), the mean RWC of *E. tef* samples dropped to 32.7% while *E. nindensis* was relatively stable at 67.6%. The mean RWC of *E. tef* samples dropped below the lethal limit to 16.3% by 8ZT on day 2 (D2), and RWC in *E. nindensis* was similarly low at 15.2% and remained relatively unchanged on D3 (14.3%). In a separate rehydration experiment, *E. nindensis* plants were desiccated to a RWC of 14.6% and re-watered (Figure 1b). *E. nindensis* recovered to a RWC of 80.8% after 12 hours of rehydration and 93.2% and 87.0% after 24 and 48 hours respectively.

**Figure 2.**
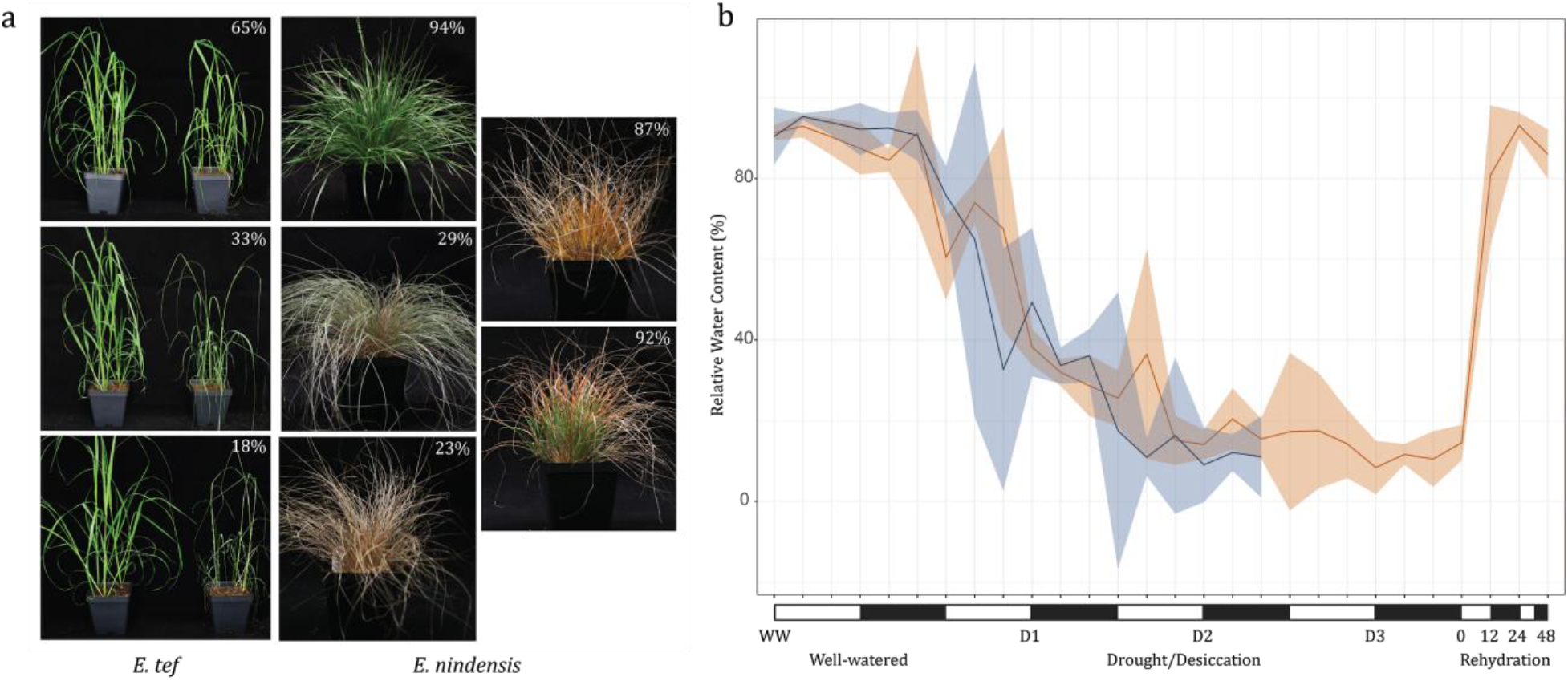
Comparative, parallel dehydration experiments in *Eragrostis* grasses. (a) Parallel drought/desiccation timecourse in *E. tef* (left) and E*. nindensis* (right). *E. nindensis* plants at 24 and 48 hours post rehydration are also shown. (b) Relative water content changes across the desiccation and rehydration experiments with 95% confidence intervals plotted for each timepoint.

In addition to RWC, we also tracked electrolyte leakage across the desiccation timecourse as a proxy for membrane damage and cell death. Electrolyte leakage was relatively low for drought D1 at 18.3% and 9.0% for *E. tef* and *E. nindensis* respectively (Supplemental Figure 4). Electrolyte leakage increased to 48.2% on D2 for *E. tef*, and from 53.0% to 64.2% for *E. nindensis* on D2 and D3 respectively. The independent rehydration experiment had lower electrolyte leakage at the desiccated timepoint (33.2%) and dropped to 11.0% and 5.2% at 12 and 24 hours post rehydration. Leaf tips and older leaves in *E. nindensis* do not always recover after desiccation, and differences in electrolyte leakage among such tissues in response to drying has been reported (Vander Willigen et al. 2001). Thus the discrepancy in electrolyte leakage between the two desiccated timepoints (D3 and R0) may be attributed to this phenomenon. The observed recovery in electrolyte leakage during rehydration suggests *E. nindensis* is able to repair desiccation induced membrane damage.

We collected leaf RNAseq data in parallel with the physiology data for eight timepoints in *E. nindensis* and three for *E. tef*. This includes control (WW) and day 1 and day 2 drought for both species (D1 and D2), as well as day 3 drought (D3), desiccated, 12 hours, 24 hours, and 48 hours post rehydration for *E. nindensi*s only (R0, R12, R24, and R48 respectively). Across the timecourse, 26,275 genes (24.4% of high confidence genes) in *E. nindensis* were differentially expressed between well-watered leaves and at least one drought or rehydration timepoint. Of these, 7,504 were upregulated and 19,506 were downregulated. Downregulated genes under drought in *E. nindensis* were significantly enriched in gene ontology (GO) terms related to photosynthesis, while upregulated genes were significantly enriched in abiotic stress response related terms, as expected (Supplemental Table 2). Similar GO term enrichment was observed for drought responses in *E. tef* (Supplemental Table 3). Many of the abiotic stress response genes that were upregulated during drought retained high levels of expression during rehydration. Genes upregulated during rehydration were significantly enriched in GO terms related to photosynthesis and specifically photoprotection and regulation of photomorphogenesis (Supplemental Table 4).

Because of the complex polyploidy of *E. nindensis* and *E. tef*, we compared expression patterns between the two species using syntenic orthogroups rather than individual gene pairs. We identified 5,600 and 6,199 syntenic groups which were upregulated during the first two water stress timepoints (D1, D2) in *E. nindensis* and *E. tef* respectively (Figure 3a). The majority of these syntenic orthogroups (3,254) were upregulated in both species (Figure 3b), supporting a broad conservation of drought responses. To further examine the degree of conservation between *E. nindensis* and *E. tef*, we compared the maximum expression in each syntenic group for the well-watered, D1, and D2 timepoints. Expression was significantly correlated between the two species (*p* > |*F*| < *0.00*) (*r*^*2*^ = *0*.*54*), suggesting that similar pathways were recruited in response to drought (Figure 3c). Nevertheless, many responses were unique to each species. Across the comparable timepoints, 2,346 syntenic orthogroups were uniquely upregulated in *E. nindensis* with no change in *E. tef* (Figure 3b). In total, 2,945 syntenic orthogroups were uniquely upregulated in *E. tef*. Candidate genes that confer vegetative desiccation tolerance are most likely to come from this set.

**Figure 3.**
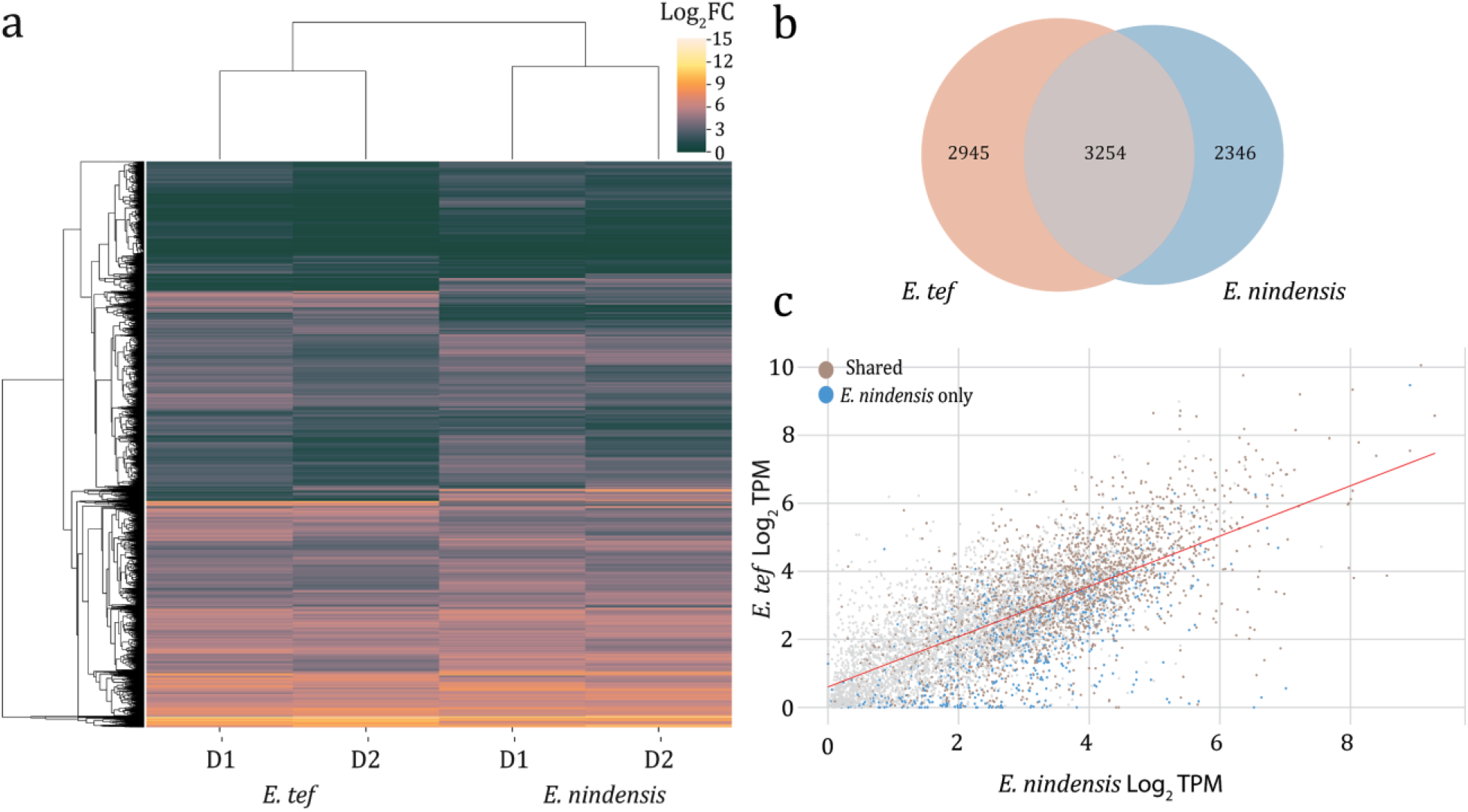
Shared and unique desiccation associated expression changes. (a) Heatmap of shared and unique syntenic orthogroups upregulated in drought/desiccation timepoints of *E. tef* and *E. nindensis.* (b) Venn diagram of shared and uniquely upregulated syntenic orthogroups. (c) Correlation of expression between the syntenic orthogroups of *E. tef* and *E. nindensis* where syntenic orthogroups with similar upregulation are shown in brown, and uniquely upregulated orthogroups in *E. nindensis* are shown in blue.

Desiccation tolerance evolved multiple times in Chloridoideae and comparative expression datasets in *E. nindensis* and *O. thomaeum* allowed us to search for patterns of convergence in these independent lineages. We compared all syntenic groups which were upregulated during any water stress timepoint between *E. nindensis*, *E. tef*, and *O. thomaeum.* We identified a set of 239 syntenic orthogroups which were upregulated in both *E. nindensis* and *O. thomaeum* but not in *E. tef*. This set of syntenic orthogroups contains many genes with seed-related functions including the seed specific vicinal oxygen chelate metalloenzyme (AT1G07645). This gene was previously shown to be upregulated in desiccated leaves and roots of the monocot resurrection plant *Xerophyta humilis* (Mulako et al. 2008). Three out of four orthologs of AT1G07645 in *E. nindensis* are upregulated between 9.8 and 11.9 fold during all of the water stress timepoints collected. Among the conserved desiccation associated genes in *E. nindensis* and *O. thomaeum* are a wide range of transcription factors with putative roles in stress or seed related functions (Table 1). Together, this suggests similar transcriptional machinery was recruited independently in multiple desiccation tolerant grass lineages.

**Table 1.**
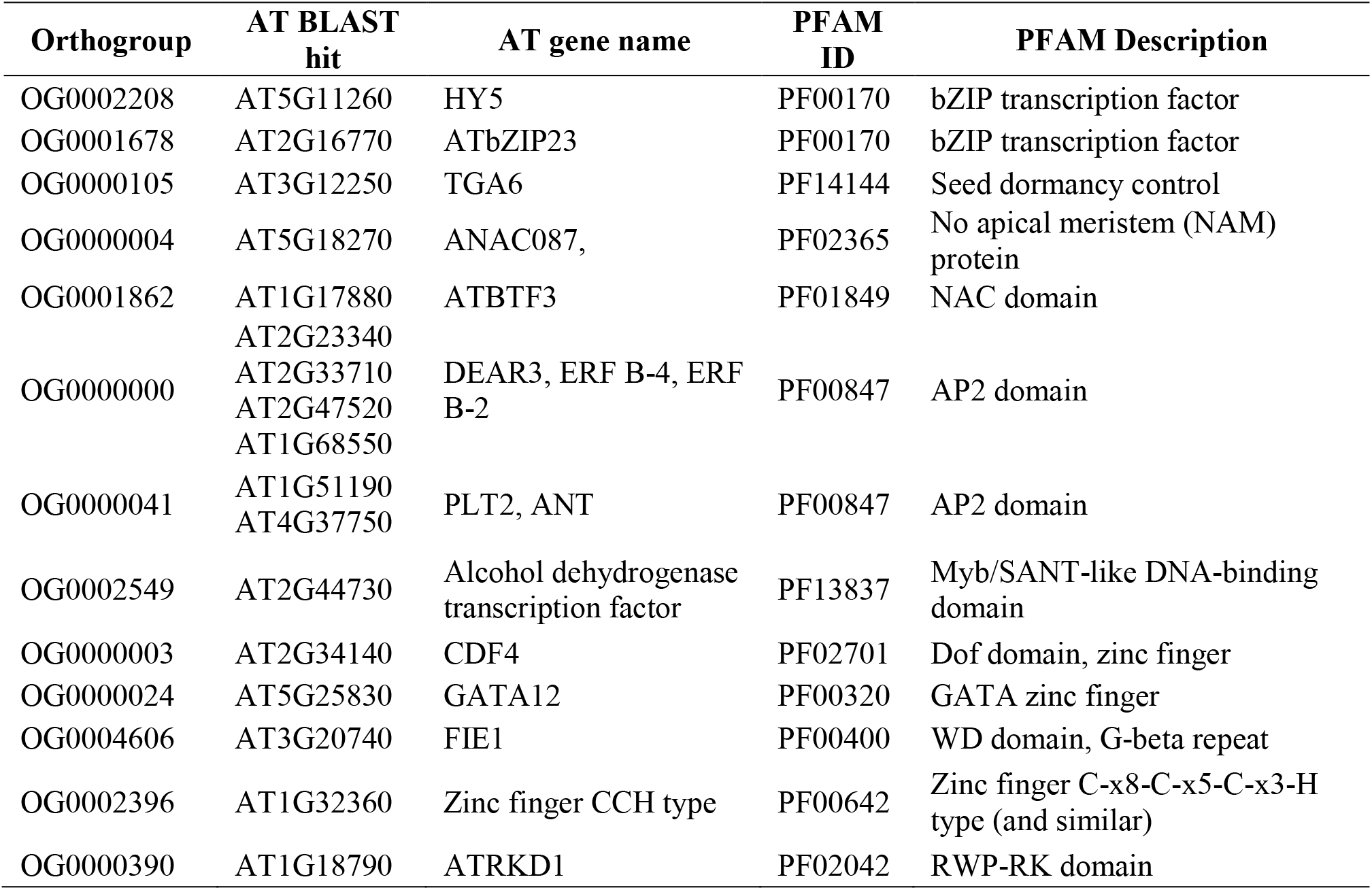
Transcription factors with desiccation specific upregulation in *E. nindensis*

### Chromatin dynamics and epigenetic changes during desiccation

Alterations of histone modifications and DNA methylation are correlated with stress induced gene expression and these processes are integral for many stress responses (Kim et al., 2015). It was previously suggested that chromatin modifications may be partly responsible for gene regulatory changes required for desiccation tolerance (Hilhorst et al., 2018; Mitra et al., 2013). We surveyed methylation changes and chromatin dynamics in well-watered and desiccated *E. nindensis* leaves using Bisulfite-seq and ChIPseq with a histone modification associated with open chromatin (H3K4me3). H3K4me3 is correlated with active transcription and these marks accumulate immediately upstream of the transcriptional start site of actively transcribed genes (Howe et al., 2017). We identified regions with differential binding of H3K4me3 antibody between well-watered *E. nindensis* leaves and desiccated leaves from the D3 timepoint. Across the three replicates of well-watered leaves, we identified 25,754 H3K4me3 peaks with significant enriched coverage (Wald test q < 0.05) over the input control (Figure 4a). The D3 samples contained 47,312 peaks and 15,832 peaks overlap by at least one base with the well-watered peaks. Despite the large number of unique peaks in each condition, only 3,757 peaks had significantly greater binding in D3 compared to WW (wald test q < 0.05), and 949 peaks had significantly more binding in WW compared to D3 (Figure 4c). We identified the closest gene to each of these differentially bound peaks and tested for enrichment of genes with up or down regulated expression in D3 compared with WW. We found significant enrichment of both up and down regulated genes among genes proximal to the peaks with increased binding in D3 (Figure 4b). Only genes downregulated in D3 compared to WW were enriched for H3K4me3 peaks. The significant number of altered H3K4me3 histone modifications and overlap with differentially expressed genes suggests chromatin dynamics play a central role in desiccation tolerance.

**Figure 4.**
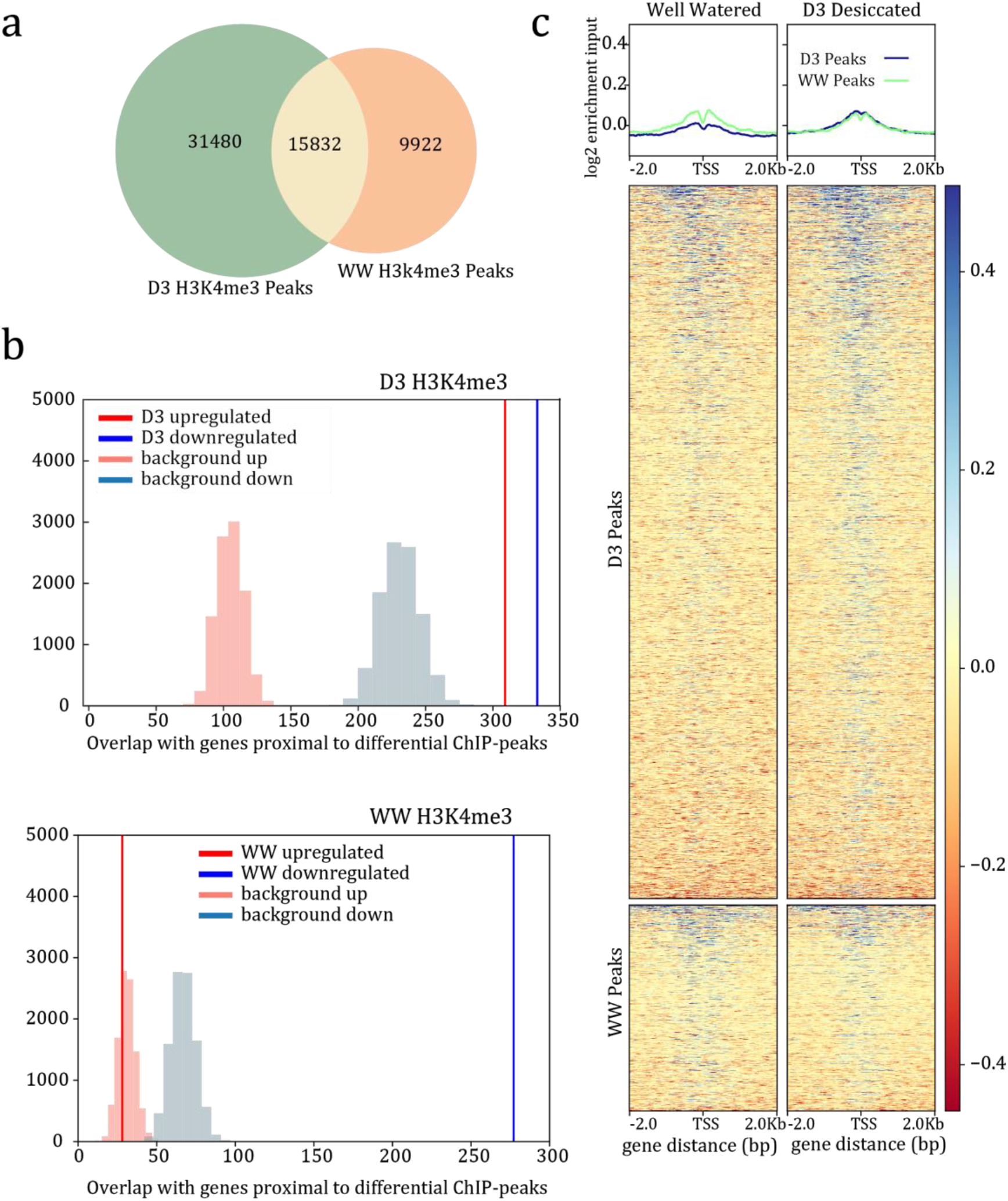
Changes in histone modifications associated with desiccation in *E. nindensis*. (a) Venn diagram of overlapping H3K4me3 peaks between well-watered and desiccated samples. (b) Enrichment of H3K4me3 peaks and upregulated genes during desiccation. The expected background distribution is shown in light blue or light red and the observed is shown in red and blue for upregulated and downregulated genes respectively. (c) Heatmap of fold enrichment of peaks at the transcriptional start site of genes. The color scale represents log2 enrichment values over the input.

We surveyed changes in DNA methylation in desiccated (D3), rehydrated, and well-watered leaf tissue. Similar to other plants, *E. nindensis* has low levels of CHH methylation across the genome, and moderate levels of CpG and CHG methylation (Figure 5a). There was no global difference in methylation levels across the surveyed drought and rehydration timepoints. However, methylation levels upstream, downstream, and within the gene body varied across the three timepoints (Figure 5b). CpG and CHH gene body methylation was significantly lower in desiccated and rehydrated leaf tissue compared to well watered. Interestingly, CHG gene body methylation was higher in rehydrated leaf tissue compared to well-watered and desiccated, with no reduction in methylation around the upstream transcriptional start site or downstream transcriptional termination site (Figure 5b). This general pattern is consistent with stress induced hypomethylation and transcriptional reprogramming (Chinnusamy and Zhu, 2009).

**Figure 5.**
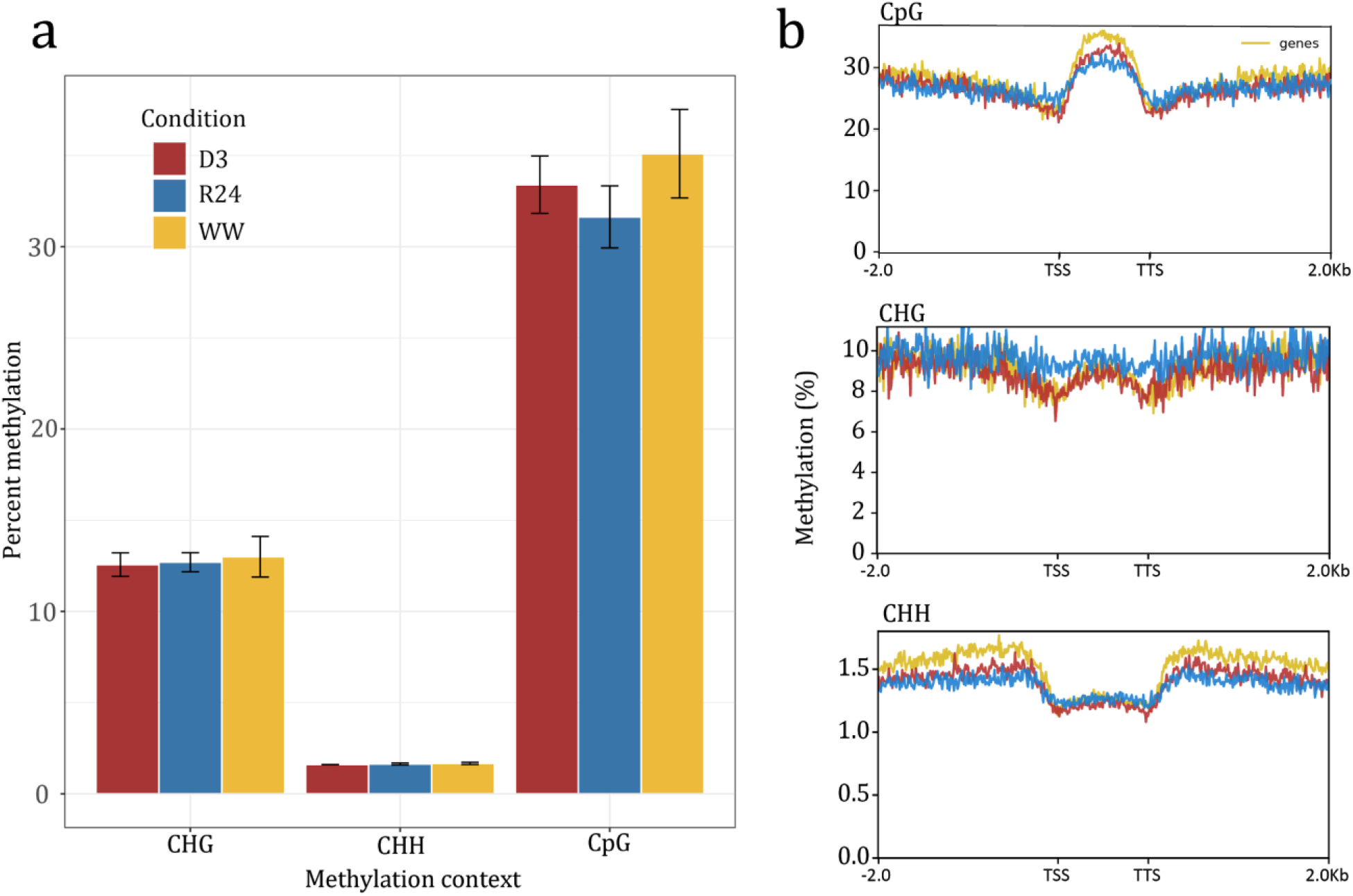
Desiccation induced changes in DNA methylation in *E. nindensis*. (a) Global patterns of CHH, CpG, and CHG methylation across the genome for well-watered (WW), desiccated (D3), and 24 hours post rehydration (R24) leaf samples. (b) Gene body methylation for the three methylation contexts in the surveyed samples. Methylation is plotted in a rolling window for all genes in the upstream (transcriptional start site; TSS), downstream (transcriptional termination site; TTS), and body of genes.

### Induction of ‘seed-related pathways’ during drought and desiccation

The longstanding hypothesis that vegetative desiccation tolerance evolved from re-wiring of seed development pathways has been supported by several recent genome-scale analyses (Costa et al., 2017; Gaff and Oliver, 2013; VanBuren et al., 2017). We used a comparative approach to test whether genes with typically seed specific expression are induced during vegetative desiccation in *E. nindensis*. We first examined gene ontology (GO) categories which were enriched among genes upregulated during drought timepoints relative to well-watered timepoints in *E. nindensis* and compared them to GO categories enriched in *E.tef* (Figure 6; Supplemental Figure 5). We classified terms into “seed related” (offspring terms of seed development GO:0048316), “stress related” (offspring terms of response to stress GO:0006950), “sugar related” (offspring terms of carbohydrate metabolic process GO:0005975), “lipid related” (offspring terms of lipid metabolic process GO:0006629), and “other” (all other terms).

**Figure 6.**
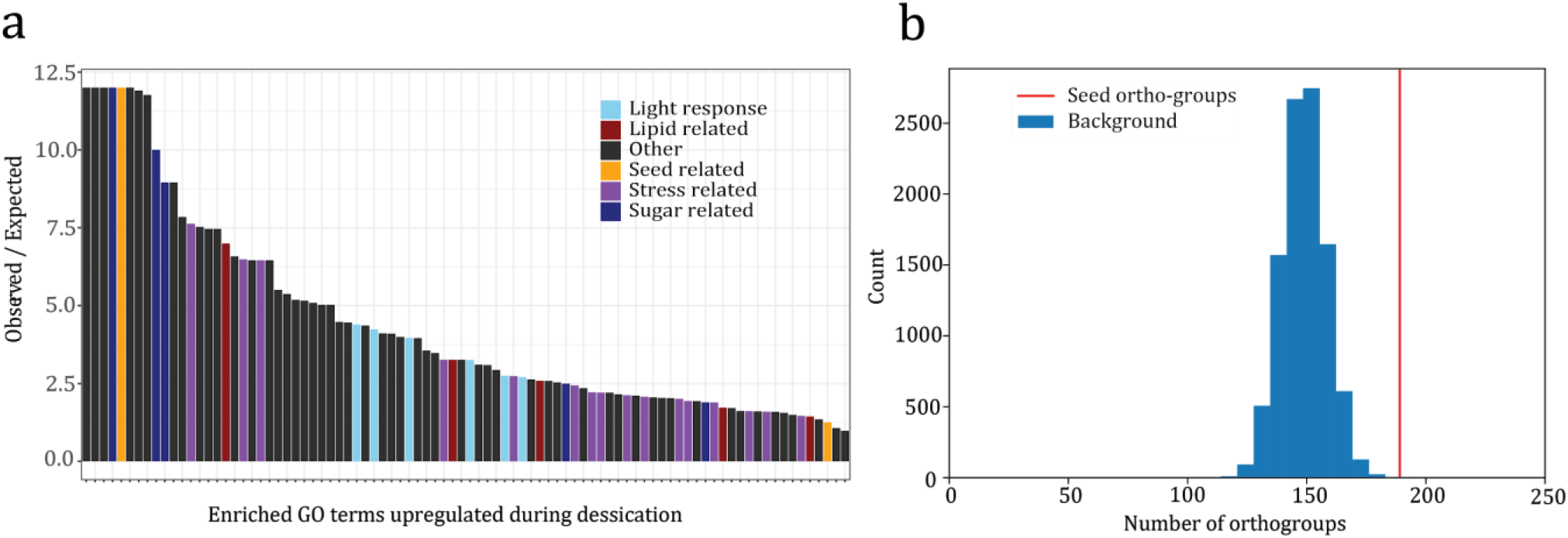
Upregulation of seed pathway genes during desiccation. (a) Enriched GO terms associated with drought/desiccation. GO terms with seed or stress related functions are highlighted. (b) Enrichment of seed related genes in *E. nindensis* (red) compared to background (blue).

To facilitate more detailed cross species comparisons, we identified groups of orthologous genes across 24 plant genomes, hereon referred to as orthogroups. We also identified pairwise syntenic orthologs between each of the six grass species with comparative expression datasets (*E. nindensis, E. tef*, *O. thomaeum, Oryza sativa*, *Zea mays*, and *Sorgum bicolor*), hereon referred to as syntelog groups (see methods). All expression comparisons were made using these orthogroups and syntegroups. We reanalyzed seed and leaf expression data from four grass species (*E. tef*, *O. sativa*, *Z. mays*, and *S. bicolor*) to create a list of “seed-related” genes with conserved expression induction in seeds relative to well-watered leaves. We then tested whether these seed related genes were overrepresented among genes upregulated in desiccating relative to well-watered *E. nindensis* leaves. A total of 189 out of 386 seed related orthogroups were upregulated in *E. nindensis* leaves in at least one drought condition (Figure 6b). Using an empirical null distribution we determined that this number was significantly more than expected by chance (*p* > |*Z*| < 0.000), suggesting that expression of seed related genes is critical for drought response in *E. nindensis* leaves.

We compared the *E. nindensis* expression data with reanalyzed well-watered, and water stressed leaf expression data from the same four grass species to determine whether the observed enrichment of seed related genes unique to desiccation tolerant plants or if it represents a more broadly shared drought response. We defined a set of seed related syntegroups such that at least one gene in each group was among the previously defined set of seed related genes with conserved upregulation in seeds relative to well-watered leaves. We then counted the number of seed related syntegroups upregulated in leaves during drought (relative to well-watered leaves) in each of the six species. *E. tef* had the greatest number of seed related genes upregulated in leaves during drought, followed by *O. thomaeum*, *E. nindensis*, and *O. sativa* (Figure 7a). In each case *S. bicolor* and *Z. mays* had fewer seed-related genes upregulated in drought treated leaves compared to the other four species. The number and severity of drought timepoints differed between species, limiting our power to draw conclusions about differences in seed expression during drought between the species. However, these data do suggest that the number of seed related genes upregulated during drought in the desiccation tolerant grasses *E. nindensis* and *O. thomaeum* is not greater than the number observed during some drought conditions in desiccation sensitive grasses.

**Figure 7.**
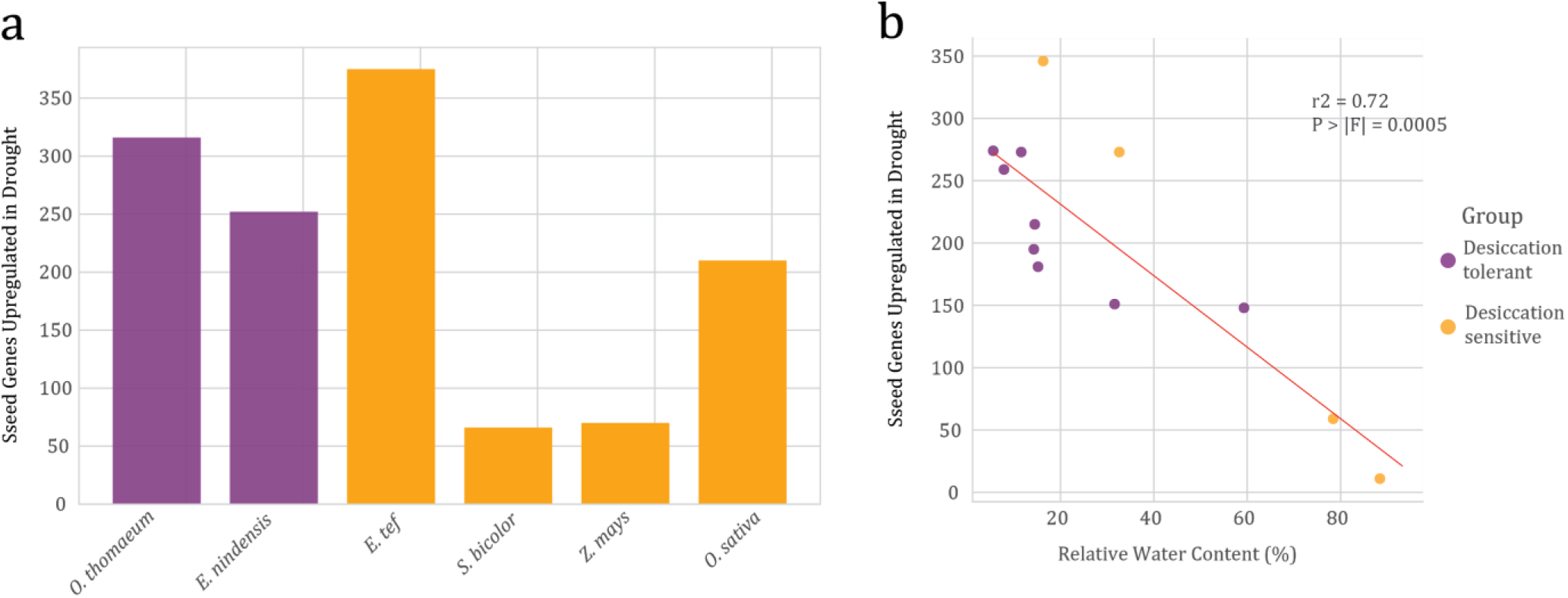
Induction of seed pathways during drought in grasses. (a) Bar graph of ‘seed-related’ gene expression in drought treated leaf tissue of the desiccation tolerant grasses *O. thomaeum* and *E. nindensis* (purple) and desiccation sensitive cereals *E. tef*, *S. bicolor*, *Z. mays*, and *O. sativa* (gold). (b) correlation of leaf relative water content (RWC) vs induced ‘seed-related’ pathways for drought treated desiccation tolerant (purple) and sensitive (gold) species.

For the four species where RWC data was available for the drought timepoints (*E. nindensis, E. tef*, *O. thomaeum*, and *S. bicolor*), we compared the number of seed related syntelog groups upregulated during each drought timepoint with the mean RWC of all replicates from that timepoint (Figure 7b). We found that RWC was a significant predictor of the number of seed related syntelogs upregulated in leaves during drought (*p* > |*F*| = 0.0005) and RWC also explained a substantial amount of the variation in the number of seed related syntelogs upregulated in leaves during drought (*r*^2^ = 0.72). Conversely, whether a sample was derived from a desiccation tolerant or sensitive plant did not significantly predict the number of seed related syntelogs upregulated in leaves during drought (*p* > |*F*| = 0.527) and this model did not explain much of the observed variation(*r*^2^ = 0.041). The spread of the model residuals was also much larger for this desiccation tolerance model as compared with the RWC model indicating a better fit for the RWC model *p* > |*F*| = 0.000503. While this analysis is somewhat limited by the confounding of species with RWC, and the paucity of both species and drought conditions included, we can conclude that more seed related genes are upregulated at lower RWC in grasses regardless of whether the species possesses desiccation tolerance.

### Induction of unique desiccation-related pathways in E. nindensis

Comparisons of drought-induced expression across grasses suggests that seed pathways are not uniquely induced in resurrection plants but instead represent a conserved response to water deficit. Using this comparative framework, we identified seed-related pathways and more general responses that are uniquely upregulated in only desiccation tolerant plants. The ABA dependent transcription factor ABI3 is important for controlling seed development and is thought to be critical for vegetative desiccation tolerance (Costa et al., 2017). We identified 23 orthogroups containing orthologs of the 98 target genes of the ABI3 regulon in Arabidopsis (Magadum et al., 2013). Of the 23 orthogroups, 16 contained at least one gene which was upregulated during drought in *E. nindensis* and 20 of the orthogroups met the same criteria in *O. thomaeum*. However, 16 of the 23 orthogroups satisfied the same criteria in *E. tef* and 12 ABI3 regulated orthogroups met those criteria in *O. sativa*. We identified one ABI3 regulated orthogroup (OG0002708) that was upregulated in leaves during dehydration in both *E. nindensis* and *O. thomaeum* but not in *E. tef* or *O. sativa*. This orthogroup (OG0002708) contains a cupin family seed storage protein (AT3G22640.1).

Late embryogenesis abundant (LEA) proteins are a group of proteins that play an important role in protecting cellular components from damage during desiccation (Goyal et al., 2005). Numerous previous works have shown that LEA genes are highly expressed during desiccation in all surveyed resurrection plants and some LEA subfamilies are expanded in the desiccation tolerant monocot *Xerophyta viscosa* (Costa et al., 2017). LEA proteins likely play an essential and conserved role in vegetative desiccation tolerance, but are also more broadly involved in drought response pathways among desiccation sensitive plants (Olvera-Carrillo et al., 2011). We identified LEA genes belonging to the 8 LEA subfamilies in the genomes of *E. nindensis*, *E. tef*, and *O. thomaeum* using PFAM domains (Supplemental Figure 6). We compared the both number and expression of each LEA subfamily between the three species (Supplemental Figure 6, Figure 8). Since each species has a different total number of annotated genes, we compared the proportion of all genes represented by each LEA subfamily rather than directly comparing the number of genes. We found no evidence of expansion of any LEA subfamilies in *E. nindensis* relative to *E. tef*. The expression of the Dehydrin, Seed Maturation Protein, LEA1, and LEA4 subfamilies was highly induced during water stress and rehydration in all three species (Figure 8). Overall, the expression of the LEA2 and LEA3 subfamilies was not induced in any of the three species, although some individual members of each family were strongly induced during water stress. Expression of LEA5 and LEA6 (also referred to as LEA18) genes was induced strongly during drought and rehydration in *E. nindensis* and *O. thomaeum* but neither subfamily showed increased expression during water stress in *E. tef.* The average transcripts per million (TPM) across all water stress conditions in *E. tef* was 1.0 and 2.6 for LEA5 and LEA6 respectively, compared to an average TPM of 7.8 and 4.9 for *E. nindensis* and 75.6 and 1,287.2 for *O. thomaeum*. Additionally, a single LEA6 gene, En_0033996, had very high expression during water stress with an average TPM of 4,211.2 across all water stress conditions.

**Figure 8.**
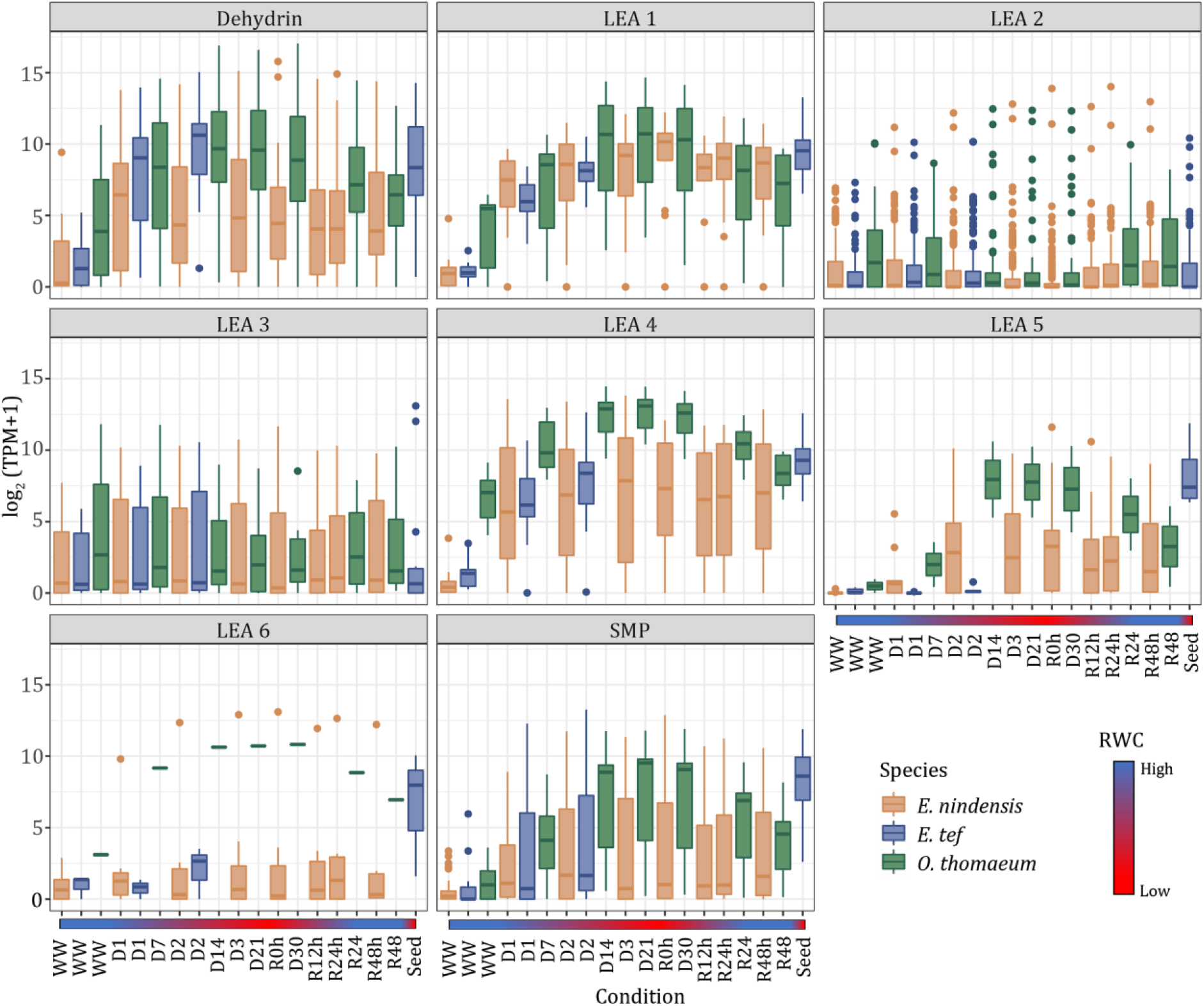
Desiccation specific induction of LEA subfamilies. The expression of the eight subfamilies of LEAs is plotted for drought and rehydration datasets from *E. nindensis*, *E. tef*, and *O. thomaeum*. Expression of each LEA gene is plotted individually with boxplots showing the distribution within a subfamily. Drying was slower in *O. thomaeum* and the D7, D14, and D21 timepoints refer to days of drought where plants were desiccated by D14.

Managing photooxidative stress is an important component of desiccation tolerance and several groups of genes related to combating excess light stress are upregulated during desiccation or rehydration in *E. nindensis*. Desiccating leaves of *E. nindensis* accumulate high levels of anthocyanins, which act as photoprotectants (Vander Willigen et al., 2001). We identified several copies of a core anthocyanin biosynthesis enzyme (UDP-glucosyl transferase) with up to 42 fold increase in expression during desiccation in *E. nindensis.* In comparison, *E. tef* showed no change in expression of UDP-glucosyl transferase enzymes during water stress. Similarly, an *E. nindensis* ortholog of the Arabidopsis tocopherol cyclase enzyme, which is critical for the production of the photoprotective carotenoid tocopherol, was upregulated during all desiccation timepoints in *E. nindensis.* The syntenic orthologs in *E. tef* were not differentially expressed under water stress. Orthologs of the ABA responsive Arabidopsis transcription factor ATAF1 (AT1G01720,) which regulates photomorphogenesis, was also upregulated during desiccation and rehydration in *E. nindensis*.

All previously sequenced resurrection plant genomes have massive tandem arrays of early light induced proteins (ELIPs) to protect against photooxidative damage during prolonged desiccation (VanBuren et al., 2019a). Consistent with this pattern, the *E. nindensis* genome has 27 ELIPs and most are found in large tandem arrays (Figure 9a). The ELIPs in *E. nindensis* are non-syntenic to the 22 orthologs in *O. thomaeum* and 5 in *E. tef*, suggesting they translocated and duplicated after the divergence of these grasses. ELIPs are induced under water stress with 23 of the 27 upregulated during desiccation or rehydration in *E. nindensis* (Figure 9b). ELIPs are most highly expressed 12 and 24 hours post rehydration, contrasting most other species where ELIPs are highest in desiccated tissues (VanBuren et al., 2019a). Unlike *O. thomaeum*, *E. nindensis* largely degrades its chlorophyll during desiccation, resulting in a comparatively slow post rehydration recovery. The high expression of ELIPs during rehydration likely protects the photosynthetic apparatus during repair, similar to patterns observed in germinating seeds (Hutin et al., 2003).

**Figure 9.**
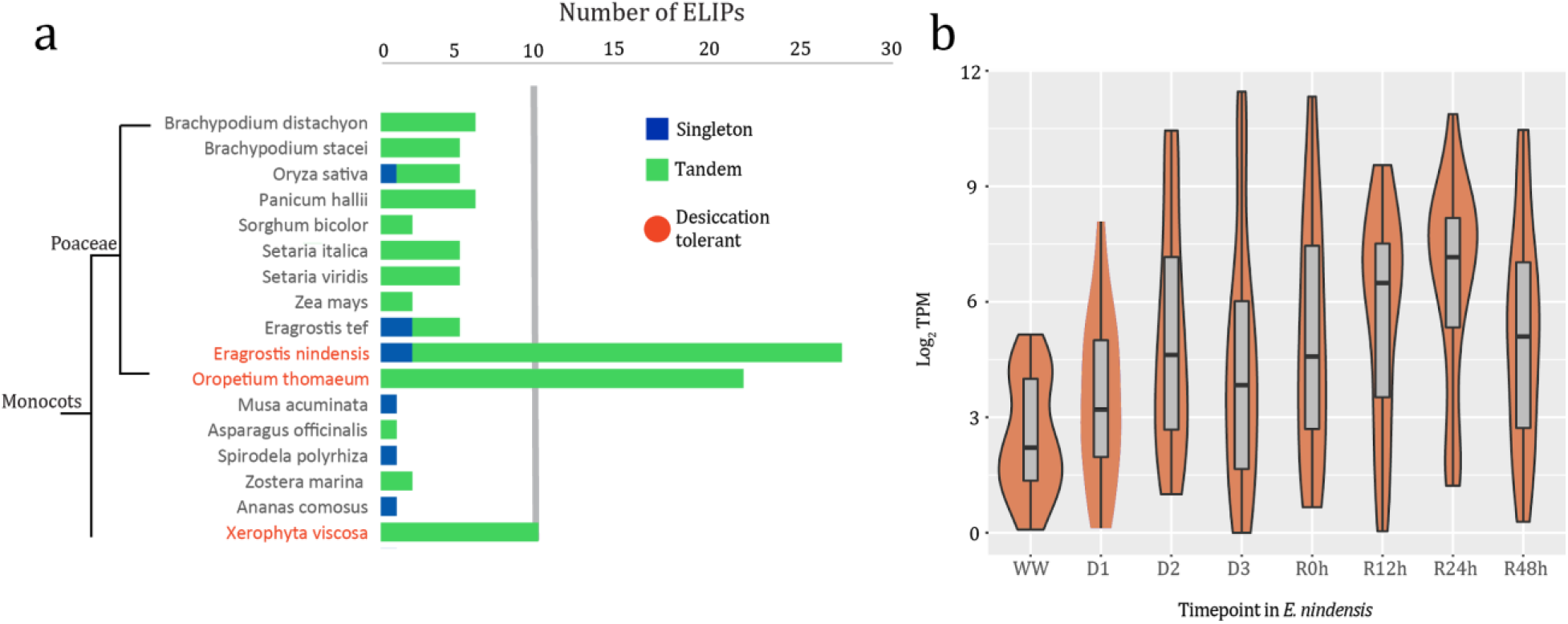
Induction of photoprotective pathways during desiccation. (a) Copy number of ELIPs in various desiccation tolerant and sensitive grasses. Tandemly duplicated ELIPs are plotted in green, and single copy or interspersed ELIPs are plotted in blue. Desiccation-tolerant species are highlighted in red. (B) Expression of ELIPs throughout desiccation and rehydration in *E. nindensis*.

The first step of chlorophyll degradation is catalyzed by chlorophyllase enzymes. Chlorophyllase genes have different expression patterns in *E. nindensis* and *O. thomaeum*, reflecting alternate strategies of chlorophyll degradation or retention during desiccation (Figure 10). An *E. nindensis* chlorophyllase (En_0076685) was upregulated 3.5 fold under desiccation but had no change in expression during any other timepoint. The syntenic ortholog in *O. thomaeum* (Ot_Chr2_06331) was not upregulated during any desiccation timepoint, but was slightly upregulated 48 hours post rehydration. The two *E. tef* chlorophyllases (Et_2A_015704 and Et_2B_019834) are upregulated in seeds but not in leaves, suggesting that desiccation associated expression of chlorophyllase may be specific to chlorophyll degrading resurrection plants. Chlorophyllases are not essential for senescence related chlorophyll degradation and a different enzyme (pheophytinase), catalyzes the removal of the phytol tail (Eckardt, 2009; Schenk et al., 2007). Pheophytinase genes are upregulated in *O. thomaeum*, *E. nindensis*, and *E. tef* under water deficit, suggesting pheophytinase activity is a more general drought response (Figure 10).

**Figure 10.**
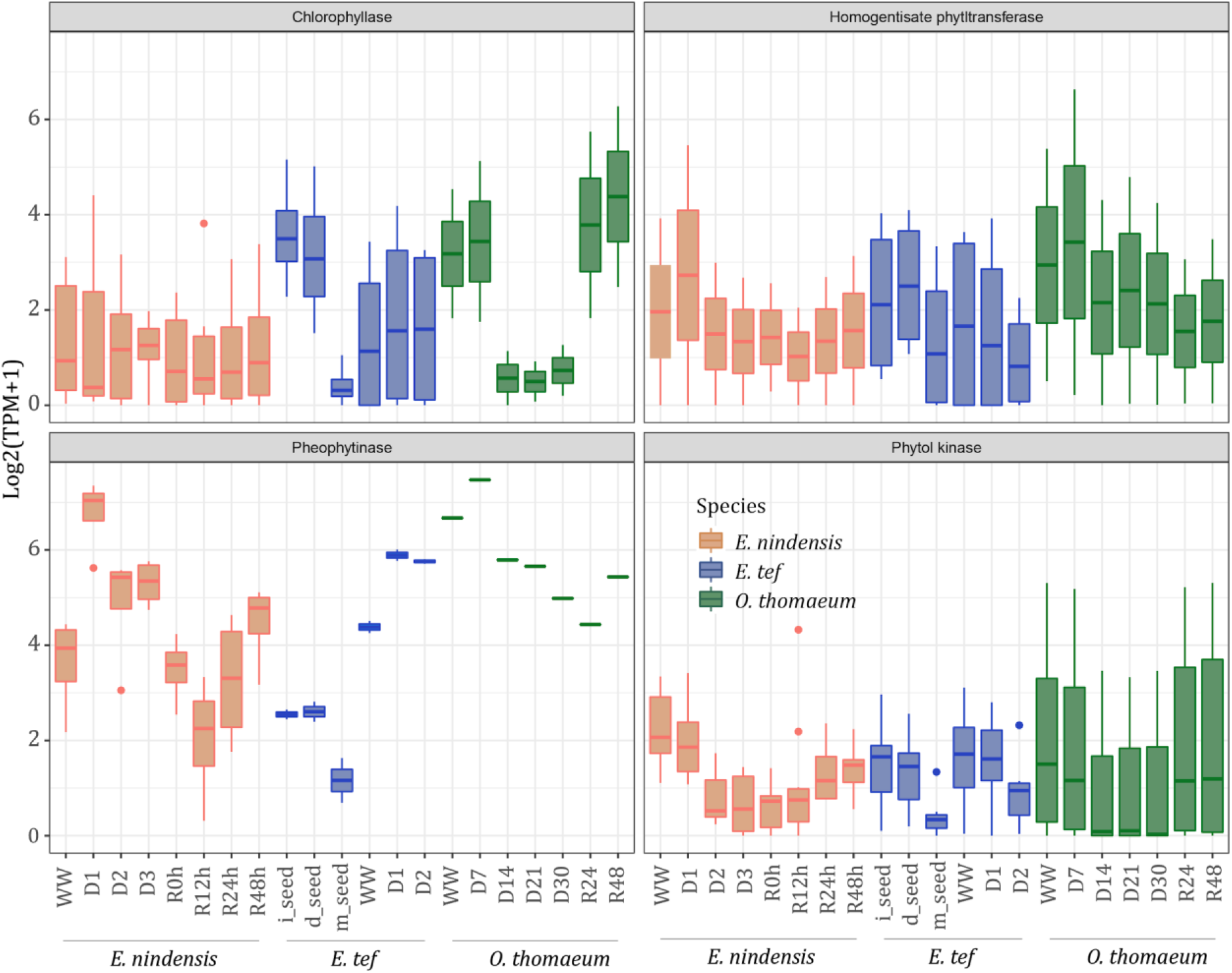
Unique desiccation associated expression of chlorophyll degradation enzymes in *E. nindensis*. The expression of several chlorophyll degradation enzymes is plotted for drought and rehydration datasets from *E. nindensis*, *E. tef*, and *O. thomaeum*. Expression of each gene is plotted individually with boxplots showing the distribution within an enzyme.

## Discussion

The Chloridoideae subfamily of grasses has at least nine independent origins of vegetative desiccation tolerance (Oliver et al., 2000). This repeated, independent evolution of vegetative desiccation tolerance has been attributed to re-wiring of seed and pollen desiccation pathways. Studies examining gene expression during desiccation have repeatedly supported this claim by identifying increased expression of seed-related genes in vegetative tissues during drying. However, few studies have conducted genome wide comparisons of desiccation tolerant species with close relative sensitive species to identify whether these seed related genes are truly unique to resurrection plants.

We observed a similar pattern of seed-related pathway expression in *E. nindenis* and the desiccation sensitive *E. tef*, suggesting these desiccation associated pathways may play a universal role in drought responses. Some seed dehydration associated pathways have well-characterized overlap with drought responses such as accumulation of osmoprotectants, LEA proteins, and ROS scavengers. More broadly, ABA-based transcription networks have some overlap in seeds and drought. For instance, 22 of the 57 LEA genes in Arabidopsis have drought induced expression, but only 10 have overlapping expression in drying seeds (Bies-Ethève et al., 2008). In *E. tef*, we observed a substantial overlap between LEA genes expressed in seeds and leaves. Many Chloridoid grasses are drought and heat tolerant and it is possible that some desiccation tolerance mechanisms are shared with desiccation sensitive but still highly resilient members this subfamily. These species may represent evolutionary intermediates with induction of some desiccation-related pathways that are not observed in less tolerant plants. Alternatively, the expression of these seed-associated genes may only occur during severe drought stress. There was a strong correlation between induction of seed-related genes and severity of the drought treatment across grass species. Most water deficit experiments are comparatively mild and few survey sublethal stresses. Seed desiccation related pathways may be induced as a last ditch effort under severe conditions, but are insufficient or too late to prevent fatal damage. Drought expression data paired with relative water content measurements was only available for two desiccation sensitive grass species. Thus, further work comparing the expression of otherwise seed specific genes during severe drought is needed to further test this hypothesis.

The ABA responsive transcription factor ABI3 is a well-characterized regulator of seed dehydration pathways. The model bryophyte *Physcomitrella patens* acquires desiccation tolerance when treated with ABA and *abi3* mutants loses this ability (Khandelwal et al., 2010). In the desiccation tolerant monocot *Xerophyta viscosa*, orthologs to the majority of of ABI3 regulated genes are expressed in leaves during dehydration (Costa et al., 2017). Similarly, we found that most of the 23 orthogroups containing ABI3 responsive genes were upregulated during desiccation in both *E. nindensis* and *O. thomaeum*. However, many of these orthogroups also had at least one gene upregulated during water stress in *E. tef* and *O. sativa*. Furthermore, expression of ABI3 orthologs was low in both *E. nindensis* and *O. thomaeum*. Thus, changes to the expression of ABI3 itself are likely insufficient to explain the vegetative desiccation tolerance phenotype. This is consistent with the fact that in Arabidopsis, both the ABA deficient mutant *aba*, and the ABA insensitive mutant *abi3*, still acquire desiccation tolerance in seeds, although the double mutant does not (Koornneef et al., 1989; Ooms et al., 1993). Drought induced expression of ABI3 regulated genes is not universal, and some are specific to desiccation tolerant tissues. Three ABI3 regulated orthogroups are expressed in both *E. nindensis* and *O. thomaeum* but not any of the sampled desiccation sensitive species, and all have seed or pollen specific expression in Arabidopsis. This supports a nuanced role of seed-related pathways where many genes overlap with typical drought responses but some crucial genes are desiccation specific in resurrection plants.

ABI5 is another major regulator of the seed dormancy and germination. Similar to ABI3, ABI5 is expressed only at trace levels in all of our samples and was not differentially expressed during any timepoint. Nevertheless, many of the downstream genes which are regulated by ABI5 are strongly upregulated during desiccation in *E. nindensis.* Furthermore, *E. nindensis* orthologs of the Arabidopsis transcription factor HY5 are induced during desiccation. HY5 was previously shown to bind to the promoter of ABI5 and regulate its expression (Chen et al., 2008). Some transcription factors have been shown to act through a “hit and run" mechanism where the transcription factor quickly dissociates from the promoters of target genes but leads to sustained transcription of the downstream targets (Para et al., 2014). If ABI3 and ABI5 act through this mechanism, it is likely that we would observe no change in expression given the limited temporal resolution of our sampling points.

Desiccation induces multiple levels of stress at the cellular level. Accumulation of LEA proteins, oil bodies, and osmoprotectants reduce stress due to cellular water loss and the resultant membrane damage. Excess light is also a major stress in desiccated leaf tissues as photooxidative damage accumulates in the absence of an active photosynthetic electron transport chain. Desiccation tolerant plants avoid photooxidative damage through either degrading chlorophyll or by protecting the chlorophyll and photosynthetic apparatus through various processes. Chlorophyll degrading species (such as *E. nindensis*) are also susceptible to photooxidative damage during rehydration when chlorophyll is re-synthesized and thylakoid damage is repaired. It was previously shown that expansion of early light induced protein (ELIP) genes is conserved among all sequenced desiccation tolerant plants (VanBuren et al., 2019a). Similar to other desiccation tolerant plants, *E. nindensis* contains an expansion of ELIPs, and the majority are highly expressed during dehydration. Interestingly, ELIPs are most highly expressed in *E. nindensis* during rehydration, contrasting patterns observed in chlorophyll retaining species. High ELIP expression was also observed during rehydration in the chlorophyll degrading species *X. viscosa (Costa et al., 2017).* ELIPs may function to protect leaves during the slow post-rehydration recovery in chlorophyll degrading species, mirroring their role in germinating seeds (Hutin et al., 2003). Consistent with adaptation to light stress, *E. nindensis* produces photoprotective pigments that are absent in *E. tef*. Furthermore, orthologs of the hub regulator for photomorphogenesis HY5 are expressed during drought exclusively in the desiccation tolerant species. Thus, we infer that mechanisms to protect against photooxidative damage are critical for the desiccation tolerance phenotype even in chlorophyll degrading species.

Here, we propose a complex but nuanced role of seed dehydration mechanisms in both drought and vegetative desiccation responses. Numerous previous studies have shows that seed dehydration pathways are important in protecting leaves of desiccation tolerant species. We identified a similar pattern in *E. nindensis*, however, comparisons to *E. tef* and other grasses reveal that seed pathways are also important in leaves of desiccation sensitive species during drought. The importance of these pathways for general drought response has been understated in the previous literature. Nevertheless, some aspects of seed dehydration pathways do appear to be specific to seeds and leaves of desiccation tolerant plants. Photoprotective pathways that are important in germinating seeds are also active in desiccated and rehydrating leaf tissue of resurrection plants. The timely, coordinated, and orderly induction of these desiccation responsive pathways may be essential for engineering improved stress resilience in crop plants.

## Methods

### Plant Growth and Sampling

Accessions of *Eragrostis nindensis* (PI 410063) and *Eragrostis tef* (PI 524434) were obtained from the USDA Germplasm Resources Information Network (www.ars-grin.gov). For the drought timecourse experiments, three seeds of *E. nindensis* were planted in 3.5" nursery pots filled with 125g of redi earth potting mix. We allowed the plants to grow for 60 days in a growth chamber under the following conditions: 12hr photoperiod, ∼ 400 mol of light, 28°C/22°C day/night temperature. Pots were brought to a total weight of 200g at the start of the drought experiment. Water was withheld for the remainder of the experiment for drought treated plants and well-watered plants were maintained at a total pot weight of 200g. Leaf tissue was sampled for relative water content (RWC) and electrolyte leakage assays every 4 hours beginning 48 hours after the start of the drought experiment. Three non-senescing leaves from each plant were randomly selected and excised at the mid-section. Samples were divided for relative water content and electrolyte leakage measurements. Inner leaves were collected for RNAseq, Bisulfite-seq and ChIP-seq experiments. Leaf samples for the D1 / WW, D2 and D3 timepoints were collected 56, 80 and 104 hours after cessation of watering respectively (8ZT on day 3, day 4 and day 5). At each timepoint, non-senescing leaf tissue was pooled from 3 plants per pot. Leaf tips were removed as they generally do not recover from desiccation. For the rehydration experiment, 102-day old plants were maintained as described above and slowly desiccated over 143 hours. Plants were then saturated with water and leaf samples were collected at 0, 12, 24 and 48 hours post rehydration. At each rehydration timepoint, samples for RWC, electrolyte leakage, RNAseq, and Methyl-seq were collected. All samples for RNAseq were flash frozen in liquid nitrogen before storing at −80°C. Samples for RWC and electolyte leakage were processed immediately after collection. Electrolyte leakage data was not collected for 48 hours post rehydration samples.

*E. tef* plants were grown in the same growth chamber as *E. nindensis* plants with the same photoperiod, light, and temperature conditions. Three *E. tef* seedlings were grown for one month in 3.5” nursery pots using redi earth potting mix. Pots were brought to the same weight at the start of the experiment (260g) and water was withheld for plants receiving the drought treatment while well-watered plants were maintained at 220g. The *E. tef* D1 and D2 samples were collected at 128 hours and 152 hours after equalizing the pot weights respectively.

### Relative Water Content

Relative water content was measured according to a previously published protocol with minor modifications (Smart, 1974). Briefly, leaf strips were excised from the midpoint of 3 *E. nindensis* leaves or a single *E. tef* leaf and immediately placed in a sealed tube at ~12°C. The fresh weight of all the samples was recorded directly following sample collection. Samples were then floated in 5mL of DI water at 4°C for 24 hours to get the turgid weight. Samples were then dried for 24-48 hours at 60°C to get dry weights. Relative water content was calculated as [(fresh weight – dry weight) / (Turgid weight – dry weight)] * 100%.

### Electrolyte leakage

Electrolyte leakage was measured according to the method outlined by A. Thalhammer (Thalhammer et al., 2014). Briefly, fresh leaf samples were placed in 5ml of deionized water and equilibrated overnight at 4°C. Samples were brought to 25°C the following day and the conductivity (conductivity_fresh_) was measured using a Mettler Toledo InLab 731-ISM conductivity probe. Samples were then boiled for 30 minutes to disrupt the cell membranes before cooling back to 25°C. The conductivity after post boiling (conductivity_boiled_) was then measured. Electrolyte leakage percentage was calculated as the (conductivity_fresh_/conductivity_boiled_) * 100%.

### Nucleic acid extraction, library preparation, and sequencing

High molecular weight genomic DNA for PacBio and Illumina library prep was isolated from leaf tissue of young *E. nindensis* plants (~30 days old) using a modified nuclei prep (Zhang et al., 1995). PacBio libraries were constructed using the manufacturer’s protocol and were size selected for 25 kb fragments on the BluePippen system (Sage Science). Libraries were sequenced on a PacBio Sequel system. An Illumina DNAseq library was constructed for polishing the PacBio based assembly using 1ug of DNA the same high molecular weight DNA prep with the KAPA HyperPrep Kit (Kapa Biosystems). The Illumina DNAseq library was sequenced on an Illumina HiSeq4000 under paired end mode (150 bp) at the RTSF Genomics Core at Michigan State University.

RNA was extracted from the timepoints described above for *E. nindensis* and *E. tef* using the Omega Biotek E.Z.N.A. Plant RNA kit according to the manufacturer’s protocol using ~200 mg of frozen tissue for each sample and quantified using the Qubit RNA HS and IQ assay kit (Invitrogen, USA). Each timepoint for RNA samples had three biological replicates. Stranded RNAseq libraries were constructed using 2ug of high-quality total RNA. The Illumina TruSeq stranded total RNA LT sample prep kit (RS-122-2401 and RS-122-2402) were used for library construction following the manufacturer’s protocol. Multiplexed RNAseq libraries were quantified, pooled, and sequenced on an Illumina HiSeq4000 under paired-end 150nt mode at the RTSF Genomics Core at Michigan State University.

### Chromatin Immuno Precipitation sequencing (ChIP-seq) library construction

Chromatin immunoprecipitation was performed according to a protocol modified from previously published protocols (Nagaki et al., 2003; Zhang et al., 2012). Briefly, nuclei were extracted from 2g of freshly ground tissue. The nuclei were then digested with micrococcal nuclease (MNase, Sigma #N5386-500UN). Following digestion a portion of the chromatin was set aside as the input control sample. The remaining chromatin was incubated overnight with a commercial H3K4me3 antibody (Abcam #Ab8580) in a rProtein A agarose (Roche #11134515001) suspension. Following antibody incubation the chromatin was eluted and purified using a DNA Clean & Concentrator kit (Zymo Research D4003). DNA-seq libraries were then constructed using the same protocol described above.

### Genome assembly

The genome size of *E. nindensis* (PI 410063) was estimated using flow cytometry in two separate runs as previously described (Arumuganathan and Earle, 1991). The *E. nindensis* genome was assembled using Canu V1.8 (Koren et al., 2017) with polishing using Pilon V1.22 (Walker et al., 2014). Raw PacBio reads were used as input for Canu and the following parameters were modified to allow for more careful unitigging and haplotype assembly: minReadLength=5000, GenomeSize=1035Mb, corOutCoverage=200 “batOptions=-dg 3 -db 3 – dr 1 -ca 500 -cp 50”. All other parameters were left as default. The output assembly graph was visualized using Bandage (Wick et al., 2015) to assess ambiguities in the graph related to repetitive elements, heterozygosity, and polyploidy. The resulting 1.96 Canu based assembly was roughly twice the estimated genome size (1.05 Gb) indicating that all four haplotypes were at least partially assembled for the allotetraploid genome. The draft Canu based contigs were polished reiteratively using Illumina paired end 150 bp data (~60x). Illumina reads were aligned to the draft contigs using bowtie2 (V2.3.0) (Langmead and Salzberg, 2012) under default parameters and the resulting bam file was used as input for Pilon. The following parameters were modified for Pilon and all others were left as default: --flank 7, --K 49, and --mindepth 10. Pilon was run recursively a total of 5 times using the updated reference for each iteration.

The *E. nindensis* genome assembly was further processed to create a pseudo-haploid representation of the genome where one of the haploypes was filtered out using the Pseudohaploid algorithm (http://github.com/schatzlab/pseudohaploid). To identify haplotype containing contigs, the genome was aligned against itself using the whole genome aligner nucmer from the MUMmer package (Delcher et al., 2003). The following parameters were used for nucmer to report all unique and repetitive alignments longer than 500 bp: nucmer -maxmatch -l 100 -c 500. This file was used as input for Pseudohaploid and the following parameters were changed in the create_pseudohaploid.sh script: MIN_IDENTITY: 95; MIN_LENGTH: 1000; MIN_CONTAIN: 90; MAX_CHAIN_GAP: 20000. Using these parameters filtered alignment chains with a minimum identity of 95%, minimum contig overlap between haplotypes of 90%, and maximum insertion size of 20kb were removed. This approach ensured that homeologous regions from the allopolyploid event were not filtered out and the strict overlap ensured that informative sequences were not purged from the assembly. The final, V2.1 assembly has a total size of 986 Mb across 4,368 contigs with an N50 of 520kb, which is similar to the expected haploid genome size.

### Genome annotation

The *E. nindensis* genome was annotated with MAKER-P v2.31.8 (Campbell et al., 2014) using transcript evidence from RNAseq data and protein homology. A de-novo transcriptome was assembled with RNAseq reads from well-watered leaf tissue using Trinity v2.6.6 (Haas et al., 2013). This assembly was used as expressed sequence tag evidence (EST) in MAKER. A second transcriptome library was assembled from RNAseq data of well-watered and desiccated leaf tissue using StringTie v1.3.3 (Pertea et al., 2015). Default parameters were used for StingTie and the ?merge option was turned on. This evidence was provided to MAKER in the “maker_gff" slot. In addition to expression evidence, protein annotations for *Arabidopsis thaliana*, *Oryza sativa*, *Sorghum bicolor, Zea mays*, *Seteria italica* and *Eragrostis tef* were used as protein homology evidence. Transposable elements and repetitive sequences were annotated using a custom repeat library (described below). We ran three rounds of ab-intio gene prediction using the SNAP gene prediction program (Korf, 2004) with the output of the prior MAKER run used as training data. BUSCO v3.0.1 (Simão et al., 2015) was used to assess the annotation quality with the set of 1440 conserved single copy orthologs from the odb9 database (https://busco.ezlab.org/v2/).

### Identification of repetitive elements

Long terminal repeat retrotransposons (LTR-RTs) were identified using LTR harvest (genome tools V1.5.8) (Ellinghaus et al., 2008) and LTR_finder V1.07 (Xu and Wang, 2007) and this list of candidate LTR-RTs were filtered and refined using LTR retriever V1.8.0 (Ou and Jiang, 2018). Parameters or LTR harvest were modified as follows based on guidelines from LTR retriever: -similar 90 –vic 10 –seed 20 –minlenltr 100 –maxlenltr 7000 –mintsd 4 –maxtsd 6. The following parameters for LTR finder were modified: -D 15000 –d 1000 –L 7000 –l 100 –p 20 –C –M 0.9. The resulting candidate LTR-RTs from both these programs were used as input for LTR retriever. LTR retriever was run with default parameters. Elements were defined as intact if they were flanked by terminal repeats. The filtered, non-redundant library from LTR retriever was used as input for whole-genome annotation of retrotransposons using RepeatMasker (http://www.repeatmasker.org/) (Tarailo-Graovac and Chen, 2009).

### ChIP-seq data analysis

Raw sequencing reads were trimmed with Trimmomatic v0.38 and aligned to the *E. nindensis* reference genome using bwa mem v 0.7.17 with default parameters (Li, 2013; Li and Durbin, 2009). Peaks of enriched ChIP signal were called relative to the corresponding input control, which was digested by MNase but not incubated with the antibody using PePr (Zhang et al., 2014). PePr accounts for the variance between replicates when calling peaks and only returns peaks that are significant after accounting for this variation. PePr was also used to identify differentially bound regions between well-watered (WW) and D3 samples. pyBedtools was used to identify the closest gene to each of these peaks including genes that overlapped the peak regions (Dale et al., 2011). The log2fold enrichment of read coverage was calculated across the genome using 10bp bins for each ChIP sample compared with the corresponding input using the bamCompare tool from deepTools v. 3.2.1 (Ramírez et al., 2014). The average log2Fold change was calculated across all three replicates of WW and D3 samples separately using WiggleTools (Zerbino et al., 2014). This average log2Fold change was plotted for the 2kb upstream and downstream regions of the transcriptional start site of the genes closest to the differential peaks using the computeMatrix and plotHeatmap tools from deepTools (Ramírez et al., 2014).

### Bisulfite-seq data analysis

Bisulfite sequencing reads were trimmed using Trimommatic v0.38 (Bolger et al., 2014). Reads were then aligned reads to bisulfite corrected *E. nindensis* reference and methylation states were called using Bismark v0.21.0 with default settings and a minimum depth of 3 reads (Krueger and Andrews, 2011). The average methylation percentage for each cytosine was calculated with a custom python script (available on gitHub). The resulting bedGraph files were then converted to bigWig format using USC genome browser’s bedGraphToBigWig script (Kent et al., 2010). The methylation percentage across gene regions was calculated with deepTools computeMatrix run in scaleRegions mode with gene bodies scaled to 1000bp and a bin size of 10bp (Ramírez et al., 2014).

### Comparative genomics

The python version of MCScan was used (https://github.com/tanghaibao/jcvi/wiki/MCscan-(Python-version)) to identify pairwise syntenic orthologs between each of the analyzed grass species (Wang et al., 2012). The *O. thomaeum* genome was used as a common anchor to the other grass genomes as it is the phylogenetically closest diploid species to *E. nindensis* and *E. tef* and it has a high quality, chromosome scale genome assembly. A minimum cutoff of five genes was used to identify syntenic gene blocks. The syntenic gene lists from all pairwise comparisons were combined and filtered into two tables, with one including syntenic orthogroups (syntegroups) with at least one gene in the three Chloridoid grasses and the second containing syntegroups with genes present in all six grass species analyzed.

While the surveyed grass genomes were largely collinear, synteny based approaches were not able to identify conserved genes that had translocated or were found in regions with extensive genome rearrangements. Orthofinder was used to identify orthologous genes that were missed by synteny based approaches. Orthofinder (v2.2.6) (Emms and Kelly, 2015) was run using default parameters with the diamond algorithm to identify orthologs in 22 species. Only orthogroups with at least one ortholog present in each species were included for the analyses.

### Expression analysis

Raw fastq files were trimmed to remove sequencing adapters using Trimmomatic v0.38 (Bolger et al., 2014). Gene expression was quantified using Salmon v0.13.1 run in quasi mapping mode (Patro et al., 2017). The transcript level estimates of expression were converted to gene level transcript per million counts using the R package tximport (Love et al., 2017). DEseq2 was used to perform differential expression analysis using the model yij ~ μ + timepoint + eij (Love et al., 2014). Each drought and rehydration timepoint was compared to well-watered to identify differentially expressed genes. The built-in wald test in the DEseq2 package was used to test whether the log2fold change of given gene was equal to 0 (Love et al., 2014). Genes with a Wald test, fdr corrected, p-value < 0.05 were considered differentially expressed.

### Identification of seed specific genes

Previously published expression data for late maturity or dry seeds and well-watered leaf tissue from four desiccation sensitive grass species (*E. tef, O. sativa S. bicolor, Z. mays*) were used to identify a set of conserved seed-related genes in grasses. All RNAseq data was downloaded from the Short Read Archive from NCBI. Seed data was reanalyzed from the following sources: *S. bicolor* (BioProject: PRJDB3281 (Makita et al. 2015), *E. tef* (VanBuren et al. 2019), *Z. mays* (Sekhon et al. 2011). The raw RNAseq data was quality filtered and quantified using the same pipeline described above. Differential expression analysis was performed using DEseq2 separately for each species in order to identify genes upregulated in seeds compared with well-watered leaves. Using this approach, 640 syntelog groups with conserved upregulation in seeds were identified among all four grasses. This list of 640 syntelog groups clustered into 386 orthogroups and was used as our list of ‘seed related’ genes. An empirical approach was used to test if these orthogroups were overrepresented among upregulated genes during desiccation in *E. nindensis.* The empirical null distribution was simulated by randomly selecting (without replacement) 386 orthogroups from the set of 11,905 orthogroups not related to seed processes. A Z-score was calculated based on the observed overlap between upregulated genes and seed orthogroups and the null distribution. This was compared to a normal distribution to determine the probability of identifying at least the observed number of genes as overlapping between the sets. Leaf drought datasets were downloaded from the NCBI SRA and analyzed as described above. The following drought datasets were analyzed: *O. sativa* (BioProject: PRJNA420056 (Fu et al. 2017), *S. bicolor* (BioProject: PRJNA319738) (Fracasso et al. 2016), *Z. mays* (BioProject: PRJNA378714).

## Data availability

The raw PacBio data, Illumina DNAseq, RNAseq data, Bisulfite-seq, and ChIP-seq are available from the National Center for Biotechnology Information Short Read Archive.. The *E. nindensis* V2.1 genome can be downloaded from NCBI, and CoGe (under ID: 54689). Code used to analyze the expression data is available on github (https://github.com/pardojer23/VanBuren_Lab_Genomics_Tools)

## Acknowledgements

We thank Yao Cao for help with DAPI staining and karyotyping and Alan Yocca for assistance with his Ka/Ks pipeline. This work is supported by funding from the National Science Foundation (MCB‐1817347 to R.V.). This publication was made possible by a predoctoral training award to Jeremy Pardo from the National Institute of General Medical Sciences of the National Institutes of Health (T32-GM110523). Hannah Chay was supported by the High School Honors Science, Math and Engineering Program at MSU.

## References

Arumuganathan K, Earle ED. 1991. Estimation of nuclear DNA content of plants by flow cytometry. Plant Mol Biol Rep 9:229–241.

Bateman RM, Crane PR, DiMichele WA, Kenrick PR, Rowe NP, Speck T, Stein WE. 1998. EARLY EVOLUTION OF LAND PLANTS: Phylogeny, Physiology, and Ecology of the Primary Terrestrial Radiation. Annu Rev Ecol Syst 29:263–292.

Bies-Ethève N, Gaubier-Comella P, Debures A, Lasserre E, Jobet E, Raynal M, Cooke R, Delseny M. 2008. Inventory, evolution and expression profiling diversity of the LEA (late embryogenesis abundant) protein gene family in Arabidopsis thaliana. Plant Mol Biol 67:107–124.

Bolger AM, Lohse M, Usadel B. 2014. Trimmomatic: a flexible trimmer for Illumina sequence data. Bioinformatics 30:2114–2120.

Campbell MS, Law M, Holt C, Stein JC, Moghe GD, Hufnagel DE, Lei J, Achawanantakun R, Jiao D, Lawrence CJ, Ware D, Shiu S-H, Childs KL, Sun Y, Jiang N, Yandell M. 2014. MAKER-P: a tool kit for the rapid creation, management, and quality control of plant genome annotations. Plant Physiol 164:513–524.

Chen H, Zhang J, Neff MM, Hong S-W, Zhang H, Deng X-W, Xiong L. 2008. Integration of light and abscisic acid signaling during seed germination and early seedling development. Proc Natl Acad Sci U S A 105:4495–4500.

Chinnusamy V, Zhu J-K. 2009. Epigenetic regulation of stress responses in plants. Curr Opin Plant Biol 12:133–139.

Costa M-CD, Artur MAS, Maia J, Jonkheer E, Derks MFL, Nijveen H, Williams B, Mundree SG, Jiménez-Gómez JM, Hesselink T, Schijlen EGWM, Ligterink W, Oliver MJ, Farrant JM, Hilhorst HWM. 2017. A footprint of desiccation tolerance in the genome of Xerophyta viscosa. Nat Plants 3:17038.

Dale RK, Pedersen BS, Quinlan AR. 2011. Pybedtools: a flexible Python library for manipulating genomic datasets and annotations. Bioinformatics 27:3423–3424.

Delcher AL, Salzberg SL, Phillippy AM. 2003. Using MUMmer to identify similar regions in large sequence sets. Curr Protoc Bioinformatics Chapter 10:Unit 10.3.

Delwiche CF, Cooper ED. 2015. The Evolutionary Origin of a Terrestrial Flora. Curr Biol 25:R899–910.

Eckardt NA. 2009. A new chlorophyll degradation pathway. Plant Cell 21:700.

Ellinghaus D, Kurtz S, Willhoeft U. 2008. LTRharvest, an efficient and flexible software for de novo detection of LTR retrotransposons. BMC Bioinformatics 9:18.

Emms DM, Kelly S. 2015. OrthoFinder: solving fundamental biases in whole genome comparisons dramatically improves orthogroup inference accuracy. Genome Biol 16:157.

Franchi GG, Piotto B, Nepi M, Baskin CC, Baskin JM, Pacini E. 2011. Pollen and seed desiccation tolerance in relation to degree of developmental arrest, dispersal, and survival. J Exp Bot 62:5267–5281.

Gaff DF, Oliver M. 2013. The evolution of desiccation tolerance in angiosperm plants: a rare yet common phenomenon. Funct Plant Biol 40:315–328.

Gechev TS, Benina M, Obata T, Tohge T, Sujeeth N, Minkov I, Hille J, Temanni M-R, Marriott AS, Bergström E, Thomas-Oates J, Antonio C, Mueller-Roeber B, Schippers JHM, Fernie AR, Toneva V. 2013. Molecular mechanisms of desiccation tolerance in the resurrection glacial relic Haberlea rhodopensis. Cell Mol Life Sci 70:689–709.

Goyal K, Walton LJ, Tunnacliffe A. 2005. LEA proteins prevent protein aggregation due to water stress. Biochem J 388:151–157.

Haas BJ, Papanicolaou A, Yassour M, Grabherr M, Blood PD, Bowden J, Couger MB, Eccles D, Li B, Lieber M, MacManes MD, Ott M, Orvis J, Pochet N, Strozzi F, Weeks N, Westerman R, William T, Dewey CN, Henschel R, LeDuc RD, Friedman N, Regev A. 2013. De novo transcript sequence reconstruction from RNA-seq using the Trinity platform for reference generation and analysis. Nat Protoc 8:1494–1512.

Hilhorst HWM, Costa M-CD, Farrant JM. 2018. A Footprint of Plant Desiccation Tolerance. Does It Exist? Mol Plant 11:1003–1005.

Howe FS, Fischl H, Murray SC, Mellor J. 2017. Is H3K4me3 instructive for transcription activation? Bioessays 39:1–12.

Hutin C, Nussaume L, Moise N, Moya I, Kloppstech K, Havaux M. 2003. Early light-induced proteins protect Arabidopsis from photooxidative stress. Proc Natl Acad Sci U S A 100:4921–4926.

Kent WJ, Zweig AS, Barber G, Hinrichs AS, Karolchik D. 2010. BigWig and BigBed: enabling browsing of large distributed datasets. Bioinformatics 26:2204–2207.

Khandelwal A, Cho SH, Marella H, Sakata Y, Perroud P-F, Pan A, Quatrano RS. 2010. Role of ABA and ABI3 in desiccation tolerance. Science 327:546.

Kim J-M, Sasaki T, Ueda M, Sako K, Seki M. 2015. Chromatin changes in response to drought, salinity, heat, and cold stresses in plants. Front Plant Sci 6:114.

Koornneef M, Hanhart CJ, Hilhorst HWM, Karssen CM. 1989. In vivo inhibition of seed development and reserve protein accumulation in recombinants of abscisic acid biosynthesis and responsiveness mutants in Arabidopsis thaliana. Plant Physiol 90:463–469.

Koren S, Walenz BP, Berlin K, Miller JR, Bergman NH, Phillippy AM. 2017. Canu: scalable and accurate long-read assembly via adaptive k-mer weighting and repeat separation. Genome Res 27:722–736.

Korf I. 2004. Gene finding in novel genomes. BMC Bioinformatics 5:59.

Krueger F, Andrews SR. 2011. Bismark: a flexible aligner and methylation caller for Bisulfite-Seq applications. Bioinformatics 27:1571–1572.

Langmead B, Salzberg SL. 2012. Fast gapped-read alignment with Bowtie 2. Nat Methods 9:357–359.

Li H. 2013. Aligning sequence reads, clone sequences and assembly contigs with BWA-MEM. arXiv [q-bioGN].

Li H, Durbin R. 2009. Fast and accurate short read alignment with Burrows-Wheeler transform. Bioinformatics. doi:10.1093/bioinformatics/btp324

Love MI, Huber W, Anders S. 2014. Moderated estimation of fold change and dispersion for RNA-seq data with DESeq2. Genome Biol 15:550.

Love MI, Soneson C, Robinson MD. 2017. Importing transcript abundance datasets with tximport. dim (txi inf rep $ infReps $ sample1) 1:5.

Magadum S, Banerjee U, Murugan P, Gangapur D, Ravikesavan R. 2013. Gene duplication as a major force in evolution. J Genet 92:155–161.

Manfre AJ, LaHatte GA, Climer CR, Marcotte WR Jr. 2009. Seed dehydration and the establishment of desiccation tolerance during seed maturation is altered in the Arabidopsis thaliana mutant atem6-1. Plant Cell Physiol 50:243–253.

Mitra J, Xu G, Wang B, Li M, Deng X. 2013. Understanding desiccation tolerance using the resurrection plant Boea hygrometrica as a model system. Front Plant Sci 4:446.

Nagaki K, Talbert PB, Zhong CX, Dawe RK, Henikoff S, Jiang J. 2003. Chromatin immunoprecipitation reveals that the 180-bp satellite repeat is the key functional DNA element of Arabidopsis thaliana centromeres. Genetics 163:1221–1225.

Oliver MJ, Tuba Z, Mishler BD. 2000. The evolution of vegetative desiccation tolerance in land plants. Plant Ecol 151:85–100.

Olvera-Carrillo Y, Luis Reyes J, Covarrubias AA. 2011. Late embryogenesis abundant proteins: versatile players in the plant adaptation to water limiting environments. Plant Signal Behav 6:586–589.

Ooms J, Leon-Kloosterziel KM, Bartels D, Koornneef M, Karssen CM. 1993. Acquisition of Desiccation Tolerance and Longevity in Seeds of Arabidopsis thaliana (A Comparative Study Using Abscisic Acid-Insensitive abi3 Mutants). Plant Physiol 102:1185–1191.

Ou S, Jiang N. 2018. LTR_retriever: A Highly Accurate and Sensitive Program for Identification of Long Terminal Repeat Retrotransposons. Plant Physiol 176:1410–1422.

Para A, Li Y, Marshall-Colón A, Varala K, Francoeur NJ, Moran TM, Edwards MB, Hackley C, Bargmann BOR, Birnbaum KD, McCombie WR, Krouk G, Coruzzi GM. 2014. Hit-and-run transcriptional control by bZIP1 mediates rapid nutrient signaling in Arabidopsis. Proc Natl Acad Sci U S A 111:10371–10376.

Patro R, Duggal G, Love MI, Irizarry RA, Kingsford C. 2017. Salmon provides fast and bias-aware quantification of transcript expression. Nat Methods 14:417–419.

Pertea M, Pertea GM, Antonescu CM, Chang T-C, Mendell JT, Salzberg SL. 2015. StringTie enables improved reconstruction of a transcriptome from RNA-seq reads. Nat Biotechnol 33:290–295.

Ramírez F, Dündar F, Diehl S, Grüning BA, Manke T. 2014. deepTools: a flexible platform for exploring deep-sequencing data. Nucleic Acids Res 42:W187–91.

Rodriguez MCS, Edsgärd D, Hussain SS, Alquezar D, Rasmussen M, Gilbert T, Nielsen BH, Bartels D, Mundy J. 2010. Transcriptomes of the desiccation-tolerant resurrection plant Craterostigma plantagineum. Plant J 63:212–228.

Schenk N, Schelbert S, Kanwischer M, Goldschmidt EE, Dörmann P, Hörtensteiner S. 2007. The chlorophyllases AtCLH1 and AtCLH2 are not essential for senescence-related chlorophyll breakdown in Arabidopsis thaliana. FEBS Lett 581:5517–5525.

Shinozaki K, Yamaguchi-Shinozaki K. 2007. Gene networks involved in drought stress response and tolerance. J Exp Bot 58:221–227.

Simão FA, Waterhouse RM, Ioannidis P, Kriventseva EV, Zdobnov EM. 2015. BUSCO: assessing genome assembly and annotation completeness with single-copy orthologs. Bioinformatics 31:3210–3212.

Smart RE. 1974. Rapid estimates of relative water content. Plant Physiol 53:258–260.

Tarailo-Graovac M, Chen N. 2009. Using RepeatMasker to identify repetitive elements in genomic sequences. Curr Protoc Bioinformatics 25:4–10.

Thalhammer A, Hincha DK, Zuther E. 2014. Measuring Freezing Tolerance: Electrolyte Leakage and Chlorophyll Fluorescence Assays. Humana Press, New York, NY. pp. 15–24.

VanBuren R, Man Wai C, Pardo J, Giarola V, Ambrosini S, Song X, Bartels D. 2018a. Desiccation Tolerance Evolved through Gene Duplication and Network Rewiring in Lindernia. Plant Cell 30:2943–2958.

VanBuren R, Pardo J, Man Wai C, Evans S, Bartels D. 2019a. Massive Tandem Proliferation of ELIPs Supports Convergent Evolution of Desiccation Tolerance across Land Plants. Plant Physiol 179:1040–1049.

VanBuren R, Wai CM, Keilwagen J, Pardo J. 2018b. A chromosome-scale assembly of the model desiccation tolerant grass Oropetium thomaeum. Plant Direct 2:e00096.

VanBuren R, Wai CM, Pardo J, Yocca AE, Wang X, Wang H, Chaluvadi SR, Bryant D, Edger PP, Bennetzen JL, Mockler TC, Michael TP. 2019b. Exceptional subgenome stability and functional divergence in allotetraploid teff, the primary cereal crop in Ethiopia. bioRxiv. doi:10.1101/580720

VanBuren R, Wai CM, Zhang Q, Song X, Edger PP, Bryant D, Michael TP, Mockler TC, Bartels D. 2017. Seed desiccation mechanisms co-opted for vegetative desiccation in the resurrection grass Oropetium thomaeum. Plant Cell Environ 40:2292–2306.

Vander Willigen C, Pammenter NW, Mundree S, Farrant J. 2001. Some physiological comparisons between the resurrection grass, Eragrostis nindensis, and the related desiccation-sensitive species, E. curvula. Plant Growth Regul 35:121–129.

Walker BJ, Abeel T, Shea T, Priest M, Abouelliel A, Sakthikumar S, Cuomo CA, Zeng Q, Wortman J, Young SK, Earl AM. 2014. Pilon: an integrated tool for comprehensive microbial variant detection and genome assembly improvement. PLoS One 9:e112963.

Wang Y, Tang H, Debarry JD, Tan X, Li J, Wang X, Lee T-H, Jin H, Marler B, Guo H, Kissinger JC, Paterson AH. 2012. MCScanX: a toolkit for detection and evolutionary analysis of gene synteny and collinearity. Nucleic Acids Res 40:e49.

Wick RR, Schultz MB, Zobel J, Holt KE. 2015. Bandage: interactive visualization of de novo genome assemblies. Bioinformatics 31:3350–3352.

Xu Z, Wang H. 2007. LTR_FINDER: an efficient tool for the prediction of full-length LTR retrotransposons. Nucleic Acids Res 35:W265–8.

Yobi A, Schlauch KA, Tillett RL, Yim WC, Espinoza C, Wone BWM, Cushman JC, Oliver MJ. 2017. Sporobolus stapfianus: Insights into desiccation tolerance in the resurrection grasses from linking transcriptomics to metabolomics. BMC Plant Biol 17:67.

Zerbino DR, Johnson N, Juettemann T, Wilder SP, Flicek P. 2014. WiggleTools: parallel processing of large collections of genome-wide datasets for visualization and statistical analysis. Bioinformatics 30:1008–1009.

Zhang H-B, Zhao X, Ding X, Paterson AH, Wing RA. 1995. Preparation of megabase-size DNA from plant nuclei. Plant J 7:175–184.

Zhang W, Zhang T, Wu Y, Jiang J. 2012. Genome-wide identification of regulatory DNA elements and protein-binding footprints using signatures of open chromatin in Arabidopsis. Plant Cell 24:2719–2731.

Zhang Y, Lin Y-H, Johnson TD, Rozek LS, Sartor MA. 2014. PePr: a peak-calling prioritization pipeline to identify consistent or differential peaks from replicated ChIP-Seq data. Bioinformatics 30:2568–2575.

Zhu Y, Wang B, Phillips J, Zhang Z-N, Du H, Xu T, Huang L-C, Zhang X-F, Xu G-H, Li W-L, Wang Z, Wang L, Liu Y-X, Deng X. 2015. Global Transcriptome Analysis Reveals Acclimation-Primed Processes Involved in the Acquisition of Desiccation Tolerance in Boea hygrometrica. Plant Cell Physiol 56:1429–1441.

