## Supplemental Information for "Intertwined signatures of desiccation and drought tolerance in grasses"

**Supplemental Figures and Tables**

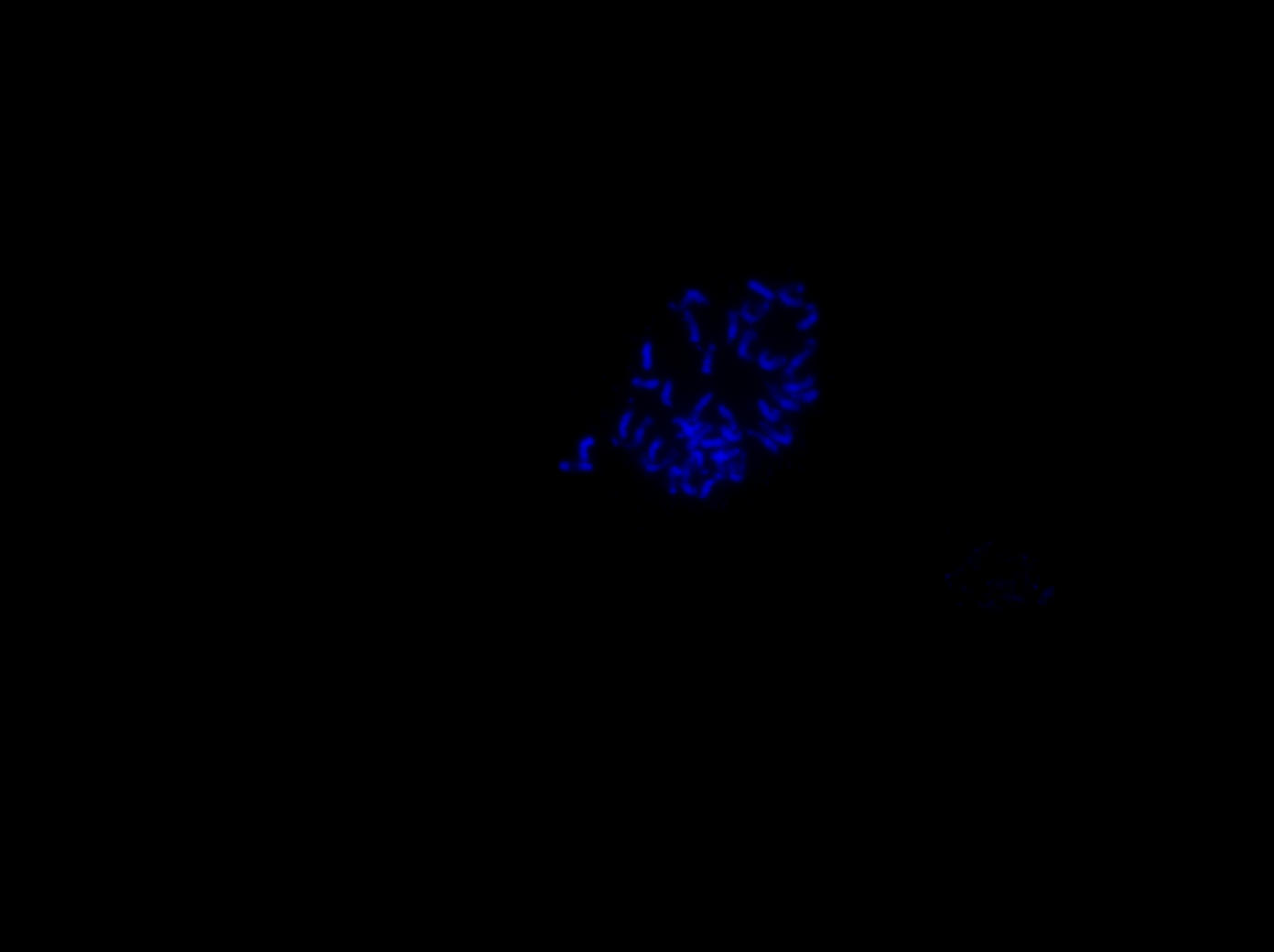

**Supplemental Figure 1. Karyotype analysis of E. nindensis. Root tip cells were stained with DAPI revealing a karyotype of 2n=4x=40.**

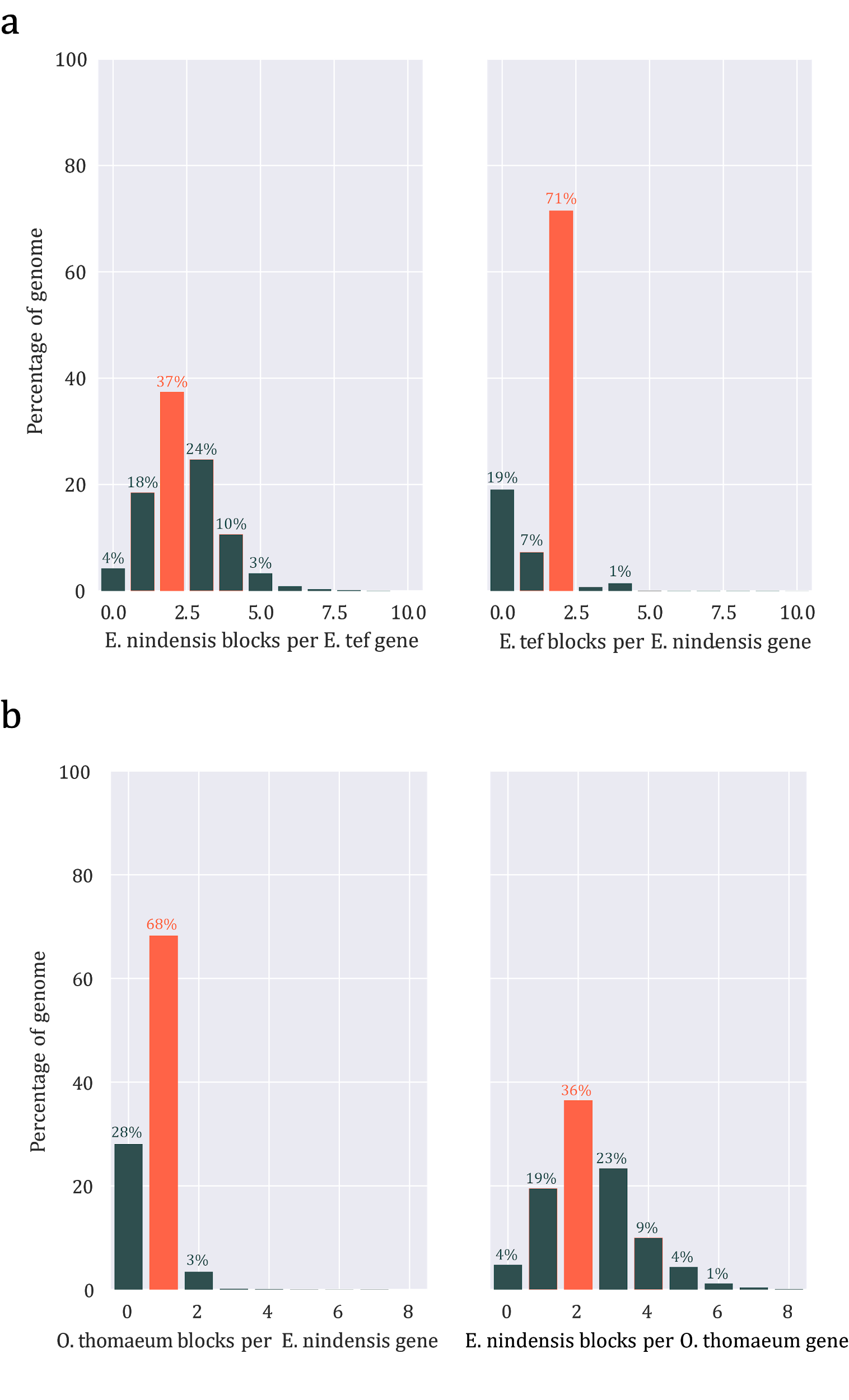

**Supplemental Figure 2. Syntenic depth of *E. nindensis* compared to (a) *E. tef* and (b) *O. thomaeum.***

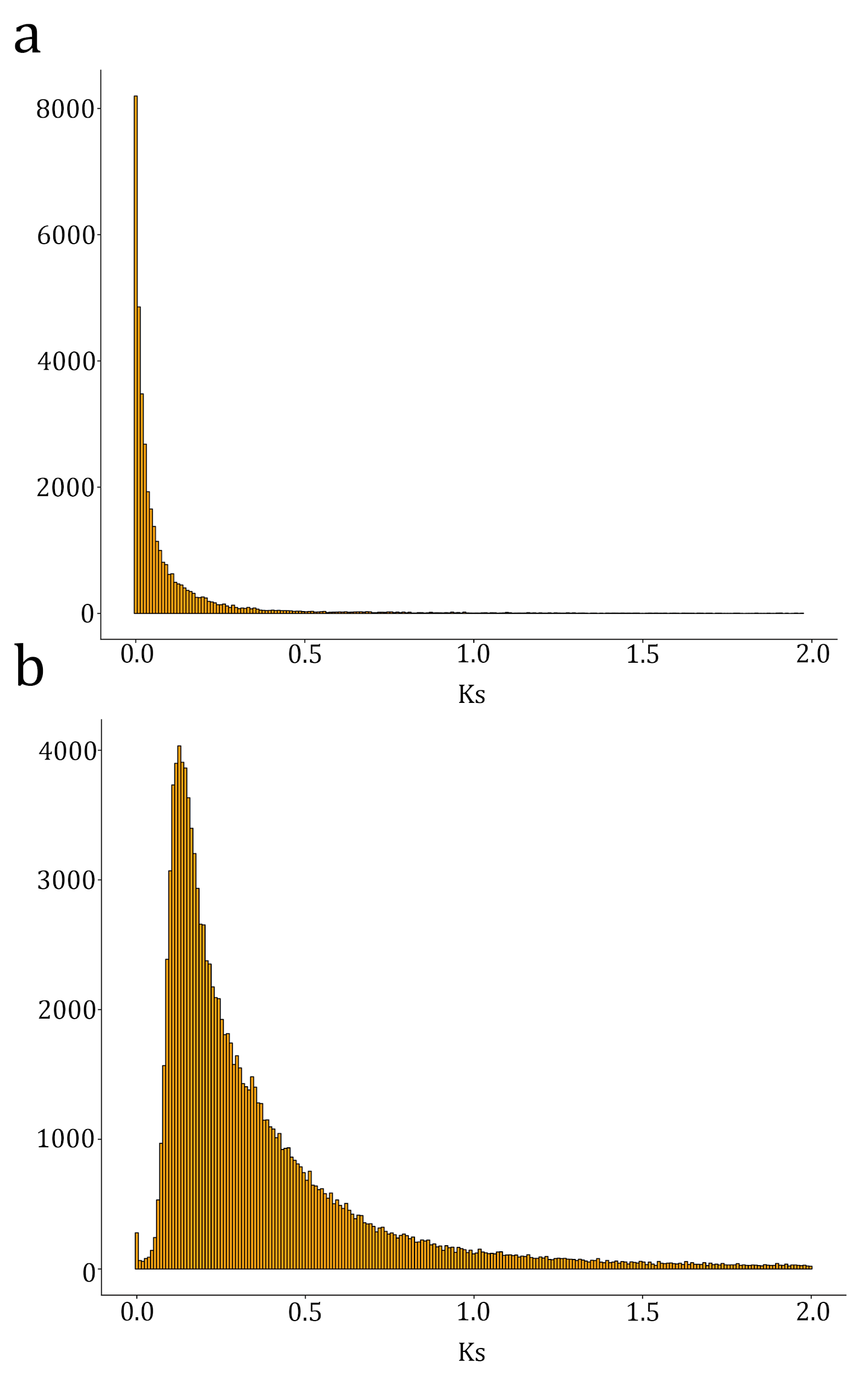

**Supplemental Figure 3 Ks distribution of duplicated genes.** (a) Ks of syntenic homeologs within the *E. nindensis* genome. (b) Ks of syntneic orthologs between the *E. tef* and *E. nindensis* genomes.

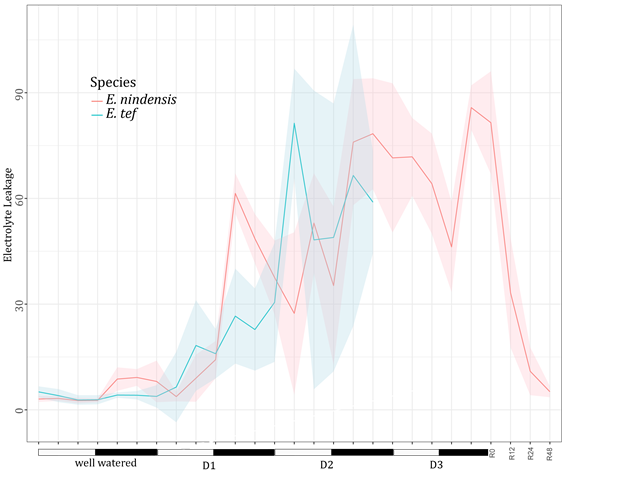

**Supplemental Figure 4. Electrolyte leakage during the desiccation and rehydration timecourse of *E. nindensis* and *E. tef*.**

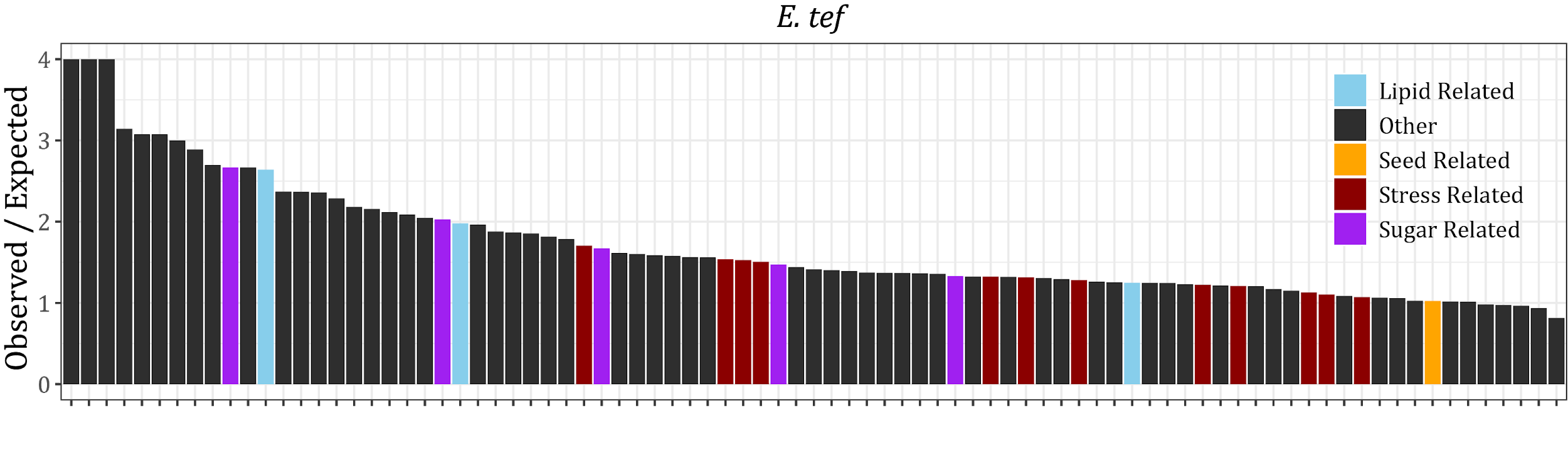

**Supplemental Figure 5. Upregulated GO terms under drought in *E. tef*.** Enriched GO terms involved in stress and seed development pathways are highlighted.

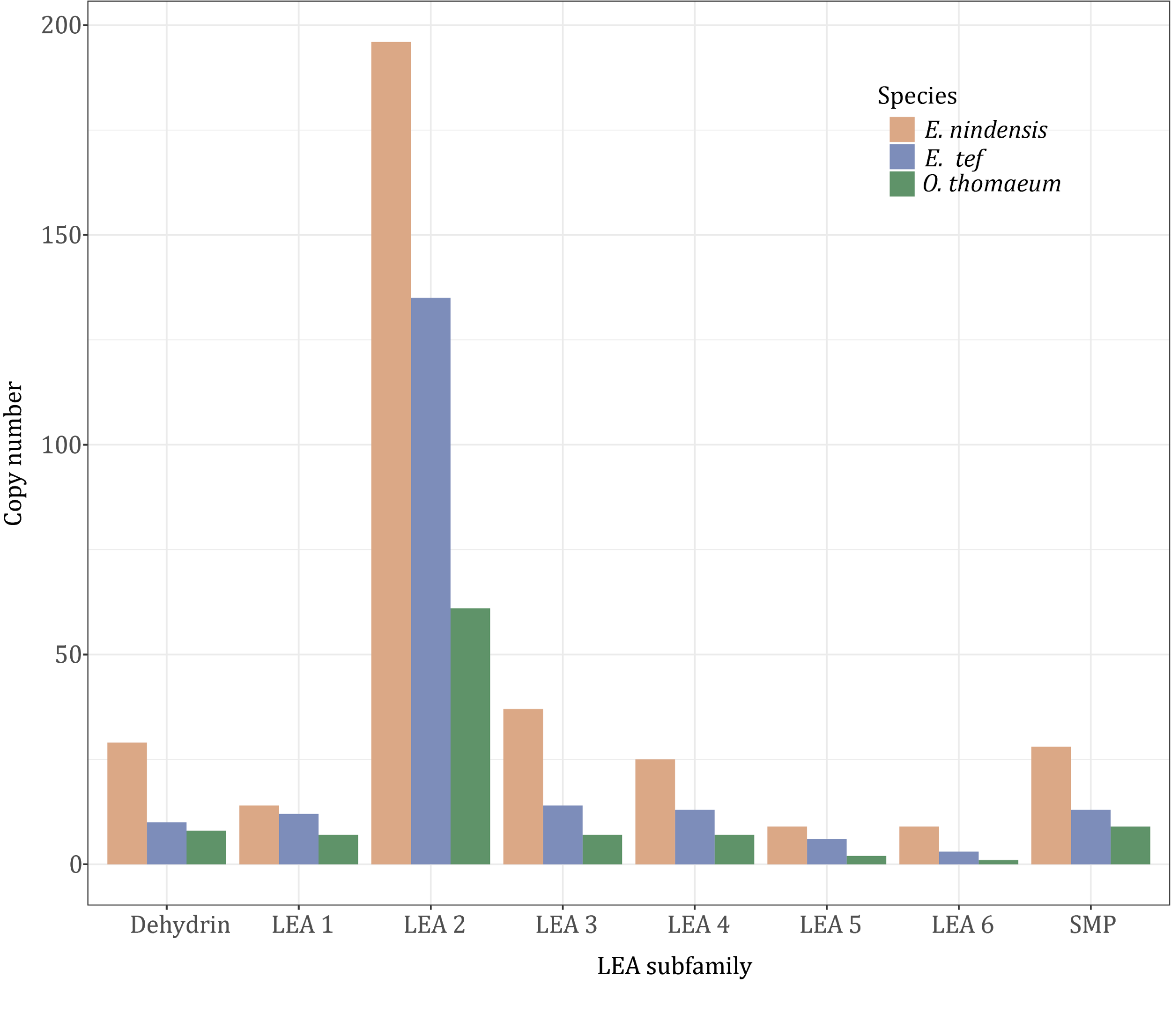

**Supplemental Figure 6. LEA family composition in Chloridoideae grasses.** The number of LEAs from each of the subfamilies is plotted for *E. nindensis*, *E. tef*, and *O. thomaeum*.

**Supplemental Table 1. Assembly statistics for the *E. nindensis* genome.**

| **Number of contigs** | **4,368** |
| --- | --- |
| **Contig N50** | **520 kb** |
| **Total length** | **986,209,651 bp** |
| **LTR composition** | **288 Mb (29%)** |
| **Number of gene models** | **116,452** |

**Supplemental Table 2. Enriched GO terms upregulated or downregulated in E*. nindensis* under the drought/desiccation timecourse. (see external csv)**

**Supplemental Table 3. Enriched GO terms upregulated or downregulated in E*. tef* under the drought timecourse. (see external csv)**

**Supplemental Table 4. Enriched GO terms upregulated or downregulated in E*. nindensis* during the rehydration timecourse. (see external csv)**
